# SpoT-mediated reduction of (p)ppGpp levels promotes *Ralstonia pseudosolanacearum* adaptation to both plant xylem and legume nodules

**DOI:** 10.64898/2026.04.03.716308

**Authors:** Nicolas Burkhardt, Mingxing Tang, Ludovic Legrand, Fabien Létisse, Philippe Vogeleer, Anthony Perrier, Alice Guidot, Delphine Capela

## Abstract

During evolution, bacteria have developed the ability to interact intimately with eukaryotic hosts. These interactions span a dynamic continuum ranging from pathogenicity to mutualism, along which bacteria can rapidly evolve and shift their lifestyles. However, the molecular mechanisms that enable bacteria to adapt to new hosts and to transition between distinct interaction modes remain poorly understood. Here, using a unique combination of two independent evolution experiments, we identified and characterized parallel adaptive mutations in *spoT*, which encodes the bifunctional (p)ppGpp synthetase-hydrolase. These mutations promote the adaptation of the plant pathogen *Ralstonia pseudosolanacearum* to two distinct plant-associated environments and two distinct lifestyles, the xylem of both susceptible and tolerant host plants as a pathogen and the root nodules of a legume as a symbiont, without compromising virulence on susceptible hosts. These mutations enhance the utilization of multiple carbon and nitrogen sources, including substrates known to be abundant in xylem sap, and increase bacterial exponential growth rate in minimal medium, suggesting reduced basal (p)ppGpp levels. Assessment of a strain deficient in SpoT synthetase activity confirmed that lowering basal (p)ppGpp levels is adaptive in both plant environments. Together, our findings reveal that fine-tuning intracellular (p)ppGpp concentrations represents an efficient strategy for optimizing bacterial adaptation to complex host-associated environments.

## Introduction

Bacteria have the remarkable ability to rapidly adapt to changing environments and colonize new ecological niches. Their adaptive evolution relies on a combination of events, including spontaneous genetic and epigenetic modifications, the acquisition of new functions through horizontal gene transfers (HGT) and the fine-tuning of gene regulatory networks (Philippe et al. 2007; Lenski 2017; Maddamsetti et al. 2017). In the context of host-bacterium interactions, there is ample evidence that bacteria can rapidly evolve and even change the nature of their relationship with their hosts (Sachs et al. 2011; Drew et al. 2021). These interactions, or symbioses, range from parasitism to mutualism on a smooth continuum, depending on the costs and benefits to the bacteria and their hosts. Shifts in interaction modes have been shaped by ecological factors, including host environmental changes and the encounter with a new host, as well as by horizontal gene transfers (HGT) of virulence or mutualistic determinants (Melnyk et al. 2019; Drew et al. 2021). For example, some plant pathogens have the propensity to expand their host range or perform host jumps (Richardson et al. 2018; Morris and Moury 2019). On the other hand, a prominent example of mutualism acquisition through HGT is the emergence of rhizobia, the nitrogen-fixing symbionts of legumes. These bacteria induce the formation of specialized organs on plant roots, called nodules, which they colonize massively and intracellularly. Within these nodules, the bacteria fix atmospheric nitrogen, converting it into ammonia, which is assimilated by the plant in exchange for carbon sources. Despite divergent outcomes, both mutualists and pathogens rely on similar molecular mechanisms to interact with their hosts (Soto et al. 2006), including protein secretion systems (mainly T3SS and T4SS) (Lomovatskaya and Romanenko 2020; Teulet et al. 2022), antimicrobial peptide transporters (LeVier et al. 2000; Smith et al. 2025) and surface polysaccharides such as exopolysaccharides, lipopolysaccharides or cyclic beta-glucans (Briones et al. 2001; Campbell et al. 2002; Soto et al. 2006).

Our research built on two long-term evolution experiments, which investigated how the same bacterium, the plant pathogen *Ralstonia pseudosolanacearum*, can evolve to either acquire the ability to infect new hosts as a pathogen or the specific ability to establish an intracellular symbiosis with legumes. The *R. pseudosolanacearum* strain GMI1000 belongs to the *Ralstonia solanacearum* species complex, which is notable for its wide host range including more than 200 plant species, its major agricultural damage, and its ability to conquer new hosts (Wicker et al. 2009; Mansfield et al. 2012; Bergsma-Vlami et al. 2018). This pathogen enters the plants through their roots, migrates to the xylem vessels where it multiplies and produces high amounts of exopolysaccharides, leading to a clogging of the vessels and the emergence of wilting symptoms characterizing the disease (Genin and Denny 2012). The first evolution experiment studied the ability of the pathogen to adapt to different host plants when propagated inside their xylem vessels (Guidot et al. 2014). The plants chosen for this experiment were either susceptible (Tomato var. Marmande, Eggplant, Pelargonium), tolerant (Cabbage, Bean) or resistant (Tomato var. Hawaii) to GMI1000. Evolved clones acquired a fitness advantage for the colonization of their host xylem after 26 to 36 successive cycles of inoculation, which corresponded to around 300 bacterial generations (Guidot et al. 2014; Gopalan-Nair et al. 2021). The second evolution experiments aimed at understanding how this pathogen can acquire the ability to become a symbiont of the legume plant *Mimosa pudica* following the horizontal acquisition of key symbiotic genes. A 557 kb symbiotic plasmid, pRalta, from the natural *M. pudica* symbiont, *Cupriavidus taiwanensis* LMG19424, was introduced into the GMI1000 strain by conjugation (Marchetti et al. 2010). This plasmid harbors a 35 kb cluster of genes essential for the establishment of symbiosis, the nodulation (*nod*) and nitrogen fixation (*nif*-*fix*) genes (Amadou et al. 2008). Following the initial selection of clones capable of nodulating *M. pudica* via spontaneous mutations, which inactivate the type III secretion system of GMI1000 (Marchetti et al. 2010), the bacteria have been submitted to successive cycles of nodulation, recovery of nodule bacteria, and reinoculation of plants (Marchetti et al. 2017). During this evolutionary process, the bacteria rapidly acquired the two first steps of symbiosis with legumes, nodulation and nodule intracellular infection, and then drastically improved these abilities (Doin de Moura et al. 2020; Doin de Moura et al. 2023).

Interestingly, some genes or pathways were parallelly targeted by mutations in both experiments, suggesting that some molecular mechanisms are adaptive in both evolutionary contexts. For example, loss-of-function mutations in components of the EfpR regulatory pathway were found to enhance the fitness of bacteria in both plant xylem and legume root nodules. The regulator EfpR controls the expression of *ca*. 200 to 900 genes. Its downregulation reduces virulence and EPS biosynthesis while enhancing metabolism and motility (Perrier et al. 2016; Capela et al. 2017).

In this study, we focused on another gene, *spoT*, that was mutated in both experiments, and that encodes the bifunctional (p)ppGpp synthetase/hydrolase. The (p)ppGpp (guanosine pentaphosphate and tetraphosphate) is a secondary messenger, also known as an alarmone, that is ubiquitous throughout the bacteria. (p)ppGpp has historically been closely associated with the stringent response, a stress response triggered by various physiological changes, such as nutrient deprivation. During the stringent response, high doses of (p)ppGpp are produced in the cells (Irving et al. 2021). These elevated levels stop bacterial growth by inhibiting many cell processes such as DNA replication, transcription, nucleotide biosynthesis, ribosomal RNA synthesis and protein translation (Irving et al. 2021). Meanwhile, genes that encode metabolic biosynthesis and nutrient transporters are induced. In *Escherichia coli*, the accumulation of (p)ppGpp, through its binding to the RNA polymerase holoenzyme, rapidly affects the expression of a large number of genes either positively or negatively, thereby having pleiotropic effects on the bacterial physiology (Sanchez-Vazquez et al. 2019). More recent research has revealed that basal levels of (p)ppGpp play significant roles beyond the high concentrations that characterize the stringent response. These basal levels have been shown to regulate growth rates in *E. coli* (Potrykus et al. 2011; Steinchen et al. 2020) and mediate the balance between stress resistance and metabolic versatility (Ferenci and Spira 2007; Zhu et al. 2019; Fernández-Coll and Cashel 2020; Spira and Ospino 2020; Steinchen et al. 2020). In most bacteria, (p)ppGpp levels are primarily controlled by a single enzyme with both synthetase and hydrolase activities. In contrast, in beta- and gamma-proteobacteria, two main enzymes encode for regulators of (p)ppGpp levels, SpoT and RelA, resulting from the duplication of the unique ancestral gene found in other bacteria (Atkinson et al. 2011). In these bacteria, SpoT keeps the two enzymatic activities, the synthesis and hydrolysis of (p)ppGpp, while RelA works solely as a synthetase (Potrykus and Cashel 2008). (p)ppGpp is synthesized by SpoT or RelA, depending on the nature of the stress (Ronneau and Hallez 2019). In response to amino acid or nitrogen deprivation, RelA acts as the primary (p)ppGpp synthetase initiating the stringent response, whereas the SpoT synthetase is activated by fatty acid starvation, carbon deprivation, iron limitation, and other stress conditions (Hauryliuk et al. 2015). However, some stress usually described as SpoT-mediated may indirectly lead to an amino acid deprivation, thereby triggering RelA-dependent (p)ppGpp synthesis, as observed in *E. coli* under fatty acid-limiting conditions (Sinha et al. 2019) or glucose deprivation (Gentry and Cashel 1996). In addition to these highly conserved enzymes, some bacterial genomes encode additional small alarmone synthetases (SAS) or small alarmone hydrolases (SAH), acting independently from the SpoT/RelA system (Irving et al. 2021). To our knowledge, the genome of our study strain, *R. pseudosolanacearum* GMI1000, as well as the symbiotic plasmid pRalta, do not possess any other gene encoding alarmone synthetases except the *relA* and *spoT* genes. However, GMI1000 does possess a potential SAH encoded by the *RSc3417* (*RS_RS17125*) gene, as well as a fourth enzyme, GppA (*RSc1537 / RS_RS07710*), involved in the conversion of pppGpp to ppGpp. These two molecules generally have a similar role, although ppGpp has a stronger overall regulatory effect than pppGpp, notably on growth rate and transcriptional activity inhibition (Mechold et al. 2013; Patacq et al. 2020).

Here, we investigated two single nucleotide polymorphisms that arose in the *spoT* gene in *R. pseudosolanacearum* experimentally evolved clones. The first mutation, *spoT*-A219P, appeared in clones evolved on cabbage (Gopalan-Nair et al. 2023), a tolerant host of GMI1000, while the second, *spoT*-L508P, emerged in clones evolved on *M. pudica* (Marchetti et al. 2017), where bacteria behave as intracellular legume symbionts. We found that both mutations increased bacterial fitness in the two distinct plant environments, the xylem of host plants and the nodules of a legume. Our results suggest that the adaptation of *R. pseudosolanacearum* to these two contrasted environments has parallelly selected for mutations that lower SpoT-dependent synthesis of basal (p)ppGpp levels.

## Results

### The SpoT mutations A219P and L508P emerged in two different evolution experiments, targeting highly conserved amino-acids

Two long-term experimental evolution studies involving the propagation of the *R. pseudosolanacearum* GMI1000 strain were previously conducted, either within the stems of six plant species, each with five independent lineages (Guidot et al. 2014; Gopalan-Nair et al. 2021), or within the root nodules of the legume *M. pudica* in 18 independent lineages (Fig. 1A; Marchetti et al. 2010; Marchetti et al. 2017; Doin de Moura et al. 2020; Doin de Moura et al. 2023). In the first experiment, five clones were randomly selected from each evolved lineage after approximately 300 bacterial generations. Five clones isolated from the lineage C after 36 passages in cabbage, a tolerant host for GMI1000, exhibited a high fitness advantage over the ancestral strain in competition assays (data from Guidot et al. 2014; Fig. 1B). The genome sequence of these clones revealed that they all carried the *spoT*-A219P mutation along with another single nucleotide polymorphism (SNP) in an intergenic region upstream from the *RSc2428* gene (Table S2). In the second evolution experiment, following the transfer of the symbiotic plasmid pRalta and an initial selection of bacteria capable of nodulating *M. pudica* (Marchetti et al. 2010), bacteria were propagated through cycles of nodulation on *M. pudica* (Marchetti et al. 2017). In one of the eighteen lineages, the lineage D, a shift in nodule intracellular infection, characterized by a higher percentage of infected area relative to the nodule section, was identified between the evolved clones D7 and D8 isolated at the 7th and 8th cycles, respectively (Fig. 1C). Among the 10 mutations that specifically occurred in the D8 clone, only the *spoT*-L508P mutation was conserved in the final evolved clone D16, isolated at the 16th cycle (Table S3) (Marchetti et al. 2017). The introduction of the *spoT*-L508P mutation into the genome of the nodulating ancestor strain (Anc^D^) or the D7 clone, led to an increase in intracellular infection of nodules to levels comparable to those observed in nodules induced by the D8 or D16 clones. Conversely, the wild-type L508 allele introduced into the D8 or D16 clone reduced their intracellular infection capacity to the same level as the ancestor (Fig. 1C).

**Figure 1.**
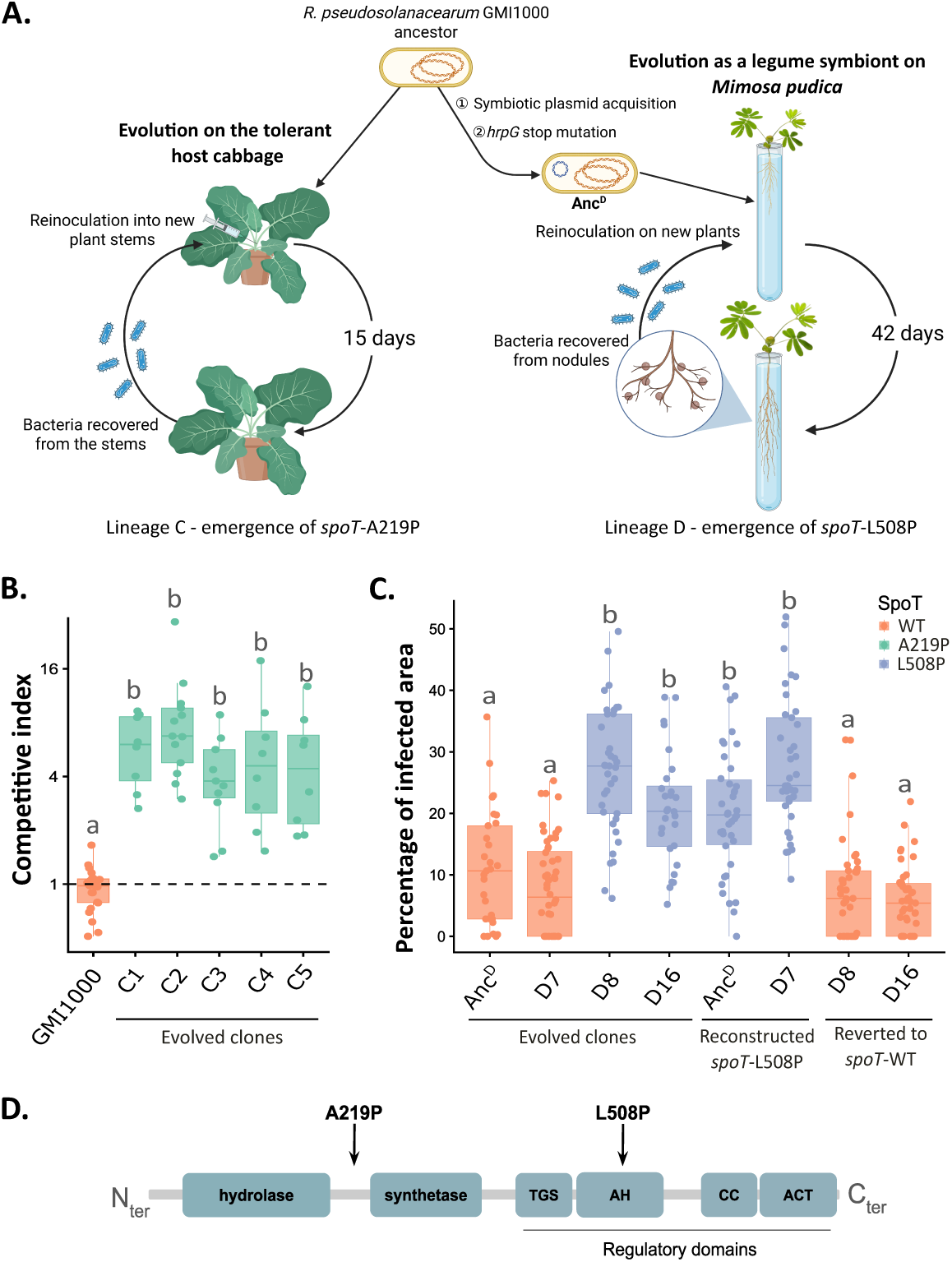
Emergence of mutations in *spoT* during the experimental evolution of *R. pseudosolanacearum* GMI1000 in plant environments. A. Schematic representation of the two independent evolution experiments initiated from the ancestral strain GMI1000. Serial passages through cabbage stems every 15 days led to the emergence of the *spoT*-A219P mutation in lineage C. In parallel, the acquisition of the symbiotic plasmid pRalta and selection of a nodulating variant mutated in *hrpG* (nodulating ancestor for the D lineage, Anc^D^) enabled serial passages through *M. pudica* nodules, leading to the emergence of the *spoT*-L508P mutation in lineage D. Created in BioRender https://BioRender.com/avhlc9y. **B. Competitive index of 5 clones evolved on cabbage** (C1-C5) carrying the *spoT*-A219P mutation after co-inoculation with the ancestral strain GMI1000 in cabbage stem. Clones C1 to C5 were isolated from the lineage C evolved population after 36 inoculation cycles on cabbage (data from Guidot et al. 2014). **C. Distribution of the percentage of infected area per nodule section** formed by clones evolved on *M. pudica*. D7, D8, D16 represent isolated clones from evolved populations of the lineage D after 7, 8 or 16 nodulation cycles on *M. pudica*. The *spoT*-L508P mutation was reconstructed in the nodulating ancestor of the D lineage (Anc^D^) and the D7 evolved clone and reverted by the wild-type allele in the D8 and D16 clones. Nodules were harvested at 21 DPI from 14 plants obtained from at least two independent experiments. Infected area data for Anc^D^ come from Capela et al. (2017). **B, C.** Different letters indicate different statistical groups (*p*<0.05, pairwise Wilcoxon test). **D. Representation of the SpoT protein** (adapted from Atkinson et al. 2011) showing the synthetase, the hydrolase, the regulatory TGS (*ThrRS, GTPase, SpoT/RelA*), AH (*Alpha-Helical*), CC (*Conserved Cysteine*) and ACT (*Aspartate kinase-Chorismate mutase-TyrA*) domains, and the positions of the A219P and L508P mutations. Raw data are available in Supplementary Table S1A. **ALT TEXT Fig. 1**: Four-panel figure showing the independent emergence of two *spoT* mutations during experimental evolution of *R. pseudosolanacearum* and the increased fitness of evolved clones carrying these mutations. Boxplots display competition assays in cabbage and percentage of infected nodule area in *M. pudica*. A protein map indicates the main functional domains of SpoT. The mutation A219P lies between the hydrolase and synthetase domains, whereas the L508P mutation is located in the AH regulatory domain.

These two amino acid residues, alanine at position 219 and leucine at position 508, are highly conserved across the phylogeny of beta- and gamma-proteobacteria according to Atkinson et al. (2011). Alanine-219 is located in an α-helical domain distinct from the known synthetase and hydrolase domains (Atkinson et al. 2011) (Fig. 1D). Leucine-508 is located within a regulatory helical domain, which may be involved in the conformational switch between the active ppGpp synthesis and hydrolysis states of the protein (Atkinson et al. 2011; Tamman et al. 2023) (Fig. 1D). Interestingly, none of these mutations have been observed in any *Ralstonia* strains sequenced to date (Fig. S1). Notably, no mutations were detected in any other enzymes known to be involved in direct (p)ppGpp metabolism in either evolution experiment (data not shown).

### The two *spoT* mutations are highly adaptive and improve bacterial proliferation in both plant xylem and legume nodules

In a first step, we assessed the adaptive character of the two *spoT* mutations in each experimental evolution context. Each mutation was reconstructed in the two ancestral backgrounds: the wild-type pathogenic ancestor GMI1000 and the minimal genetic background required for nodulation on *M. pudica,* GMI1000 pRalta *hrpG* (Marchetti et al. 2010). The *in planta* fitness of the mutants was estimated using competitive indexes (CIs) by co-inoculating the mutant strains and their ancestors, both of which tagged with different fluorophores (either mCherry or GFP). The *spoT*-A219P mutant from the cabbage-evolved lineage, exhibited an increased fitness for colonizing cabbage xylem. This increase in fitness is not specific to cabbage, as the *spoT*-A219P mutant was also more fit than the wild-type strain for colonizing the xylem of the susceptible host tomato var. Marmande (Fig. 2A). Interestingly, the *spoT*-L508P mutant, from the *M. pudica* evolved lineage, also demonstrated an increased fitness for xylem colonization in both tolerant (cabbage) and susceptible (tomato) hosts. To assess whether this fitness gain was dependent on the presence of the wild-type strain in the inoculum, we also performed single strain inoculations of tomato plants with mutant and wild-type strains (Fig. 2B). The two *spoT* mutants were not affected for their bacterial multiplication in xylem, suggesting that their adaptive character is not dependent on a cross-complementation by the wild-type strain. The *spoT*-A219P mutant even showed slightly higher bacterial proliferation than the wild-type strain in tomato xylem at 5 days post-inoculation (DPI) (Fig. 2B). In symbiosis with *M. pudica*, the two *spoT* mutations reconstructed in the nodulating genetic background also triggered a very high increase in fitness (Fig. 2C). The nodules formed by both *spoT* mutants harbored a 10-fold increase in bacterial load, revealing that the nodules are better invaded (Fig. 2D). Overall, the two mutations, *spoT*-A219P and *spoT*-L508P, were found to be highly adaptive in both evolutionary contexts, similarly leading to increased fitness in both plant xylem and legume root nodules.

**Figure 2.**
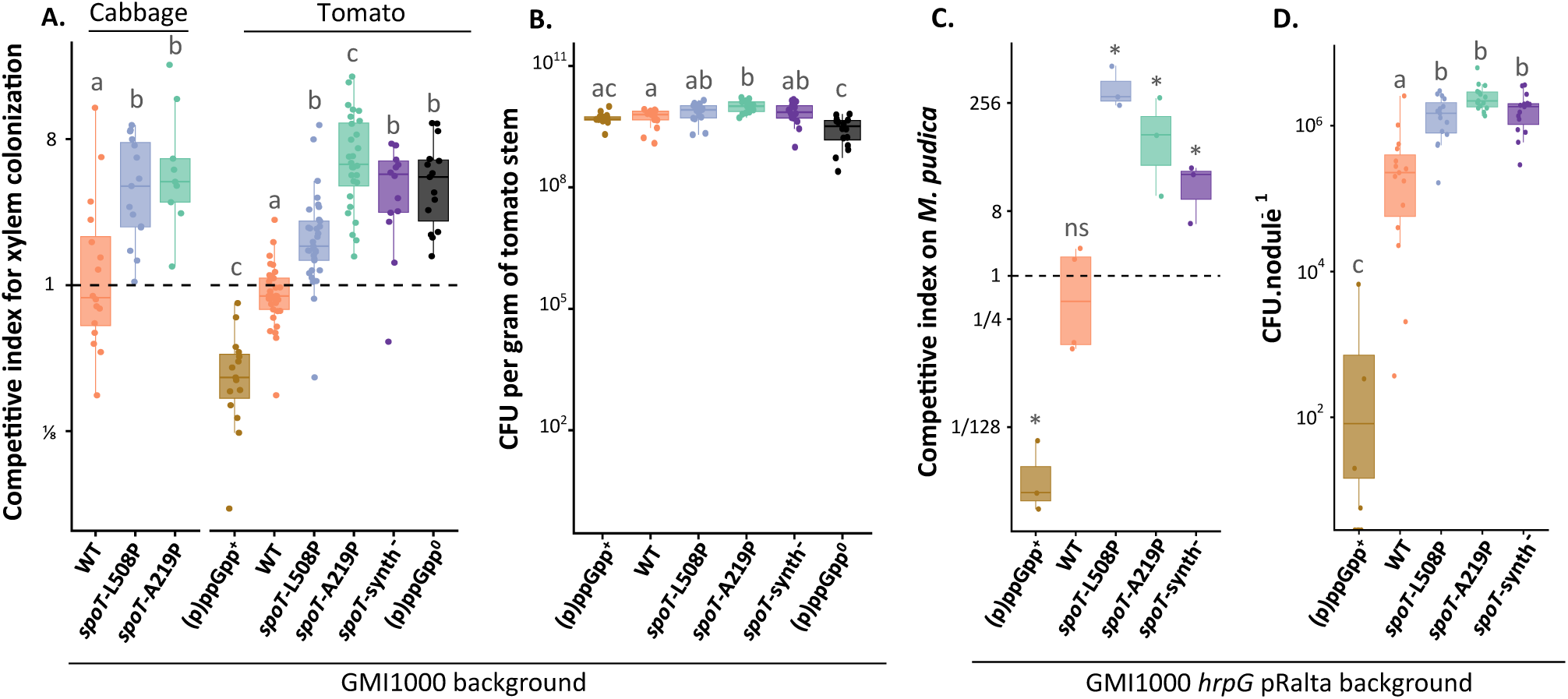
*In planta* fitness of *spoT* and (p)ppGpp synthesis mutants. **A. Competition assays for xylem colonization.** Plant stems were co-inoculated with 1:1 mixes consisting of the wild-type GMI1000 mCherry-tagged strain and a competing GFP-tagged strain (except the (p)ppGpp^+^, which is untagged). Bacteria were recovered 5 days after inoculation from the plant stems and plated. The ratios of the number of GFP-tagged colonies to the number of mCherry-tagged colonies were determined using fluorescence imaging. Each dot represents the competitive index (CI) obtained from a single plant, calculated as CI = final ratio / initial ratio. **B. Infection of tomato stems after single strain inoculations.** Bacteria were injected into the stem and counted by plating after 5 days. The number of bacteria was normalized by the weight of the stem. **C. Competition assays on *Mimosa pudica*.** *M. pudica* plants were co-inoculated with 1:1 mixes of the GMI1000 pRalta *hrpG* mCherry-tagged strain and a competing GFP-tagged strain (except the (p)ppGpp^+^, which is untagged). Bacteria were recovered 21 days after inoculation from nodules and plated. Each dot represents the competitive index obtained from a pool of nodules from 20 plants. * CI statistically different from 1, ns: not significantly different from 1 (Student *t*-test after log transformation, *p* < 0.05). **D. Infection of M. pudica nodules after single strain inoculations**. Bacteria were recovered from nodules 21 days after inoculation and counted by plating. **A, B, C, D** The (p)ppGpp^0^ strain lacks both (p)ppGpp synthetases (Δ*relA* Δ*spoT)*, whereas the (p)ppGpp^+^ strain harbors an additional chromosomal copy of the *relA* synthetase gene under the control of a constitutive promoter, and the *spoT*-synth^-^ strain refers to a *spoT*-E333Q mutation inactivating the synthetase activity of the SpoT protein. **A, B, D** Statistical analyses were performed using a pairwise Wilcoxon test. Different letters indicate significantly different conditions (*p* < 0.05). **A, B, C, D** Data are from 3-8 independent experiments with 5-20 plants per experiment. Raw data are available in Supplementary Table S1B. **ALT text Fig. 2**: Four-panel figure comparing the fitness of strains affected for (p)ppGpp synthesis in different host plants. Boxplots show the relative fitness of mutants compared to the wild-type strain in cabbage and tomato xylem (A), the bacterial loads in tomato stems after single strain inoculation (B), the relative fitness in *Mimosa pudica* nodules (C), and the bacterial load per nodule (D). Mutations in *spoT* increase fitness in both xylem and nodules. Statistical groupings are indicated.

### The pathogenicity of the *spoT* mutants is unchanged

As the *spoT-*A219P and L508P mutants were more efficient to colonize their host, we assessed their virulence on the susceptible host for GMI1000, tomato var. Marmande. We inoculated plants either through soil drenching, mimicking the usual infection process of *R. pseudosolanacearum*, or via stem injection, reflecting the experimental evolution setting. The appearance of wilting symptoms was monitored and used to calculate a hazard ratio, describing the virulence of each strain relative to the wild-type ancestor. No significant differences in virulence were observed between the *spoT* mutants and the wild-type strain when using both inoculation methods (Fig. 3). The virulence of the two *spoT* mutants is thus unchanged, suggesting that the expression of pathogenicity determinants are not affected in these strains.

**Figure 3.**
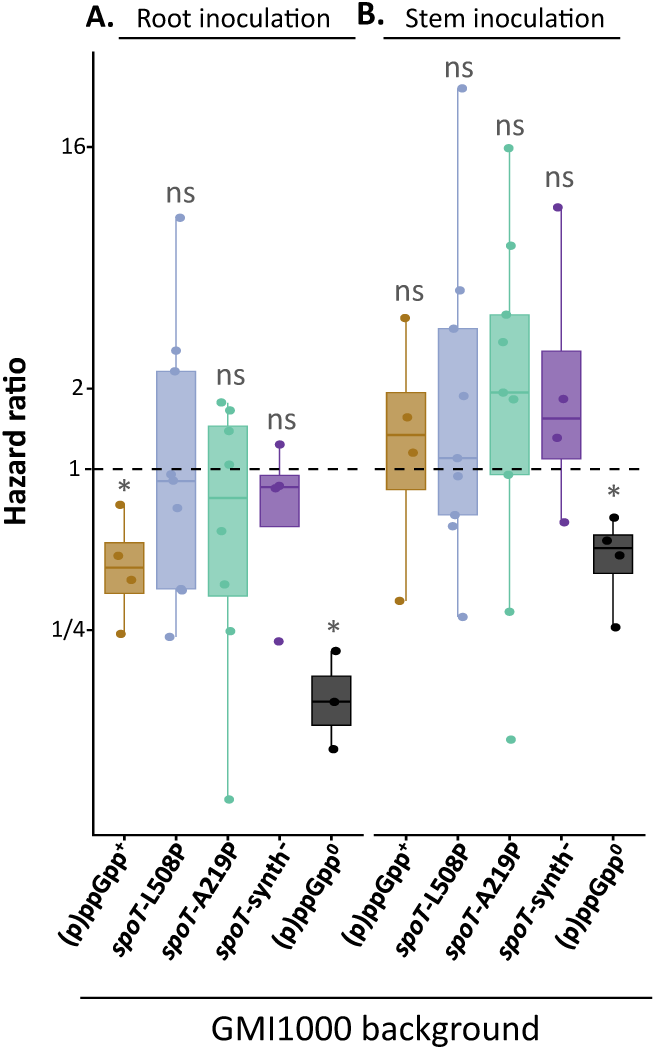
Pathogenicity assays on susceptible tomato plants. **A.** Roots were inoculated by pouring a suspension of 2.5.10^10^ bacterial cells for 16 plants. **B.** Stems were directly inoculated by injecting 10^4^ bacterial cells. Symptom progression was recorded daily. Each dot represents the hazard ratio, indicating the relative risk of symptom development for each mutant compared to the wild-type strain GMI1000, for each independent replicate (3 to 9). ns, non-significantly different from 1. * Significantly different from 1 (*p* < 0.05, t-test on values transformed by natural logarithm). Raw data are available in Supplementary Table S1C. **ALT text Fig. 3**: Two-panel figure showing hazard ratios in susceptible tomatoes following root inoculation (A) or stem injection (B). Boxplots compare strains affected for (p)ppGpp synthesis relative to the wild-type strain (dashed line = 1). Most mutants show hazard ratios that are not significantly different to the wild type, except for the (p)ppGpp^0^ strain, which displays reduced virulence with both inoculation methods, and the (p)ppGpp⁺ strain, which displays reduced virulence with root inoculation.

### The *spoT* mutations enhance the utilization of several carbon and nitrogen sources

To investigate the enhanced fitness of the *spoT* mutants in plants, we assessed their metabolic capacities using Biolog phenotype microarrays designed to test the utilization of carbon and nitrogen sources through the reduction of a tetrazolium redox dye by NADH production during substrate catabolism. Both *spoT* mutants showed increased tetrazolium reduction on several substrates compared to the wild-type strain, indicating enhanced metabolic activity (Table 1). Although the sets of substrates that were significantly better metabolized differed between the two mutants, several carbon compounds were consistently better utilized by both strains, including sugars (D-fructose, D-galactose, D-saccharic acid, D-trehalose and sucrose) and amino acids (L-alanine, L-glutamine, L-histidine, L-proline, L-serine and L-threonine). Among the nitrogen sources that were more efficiently utilized by the *spoT* mutants were adenine, xanthine, L-histidine and nitrite. Interestingly, several of the sugars and amino acids better utilized by the *spoT* mutants have been detected in tomato xylem (Table 1) (Lowe-Power et al. 2018; Baroukh et al. 2022). In particular, L-glutamine was identified as the main carbon source used by *R. pseudosolanacearum* in xylem sap (Baroukh et al. 2022). To assess whether the slight increase in utilization of this amino acid, as measured by Biolog (Table 1), indicates an enhanced ability to grow on these substrates, we measured the growth rates of the mutants in a synthetic minimal medium in which L-glutamine was the sole carbon source. In this medium, both *spoT*-A219P and *spoT*-L508P mutants displayed higher growth rates compared to the wild-type strain (+14,7% and +15,6% respectively, Fig. 4). Overall, Biolog data indicate that *spoT* mutations increase the capacity of the bacteria to utilize several carbon and nitrogen sources, which may have resulted in faster proliferation in plant environments.

**Figure 4.**
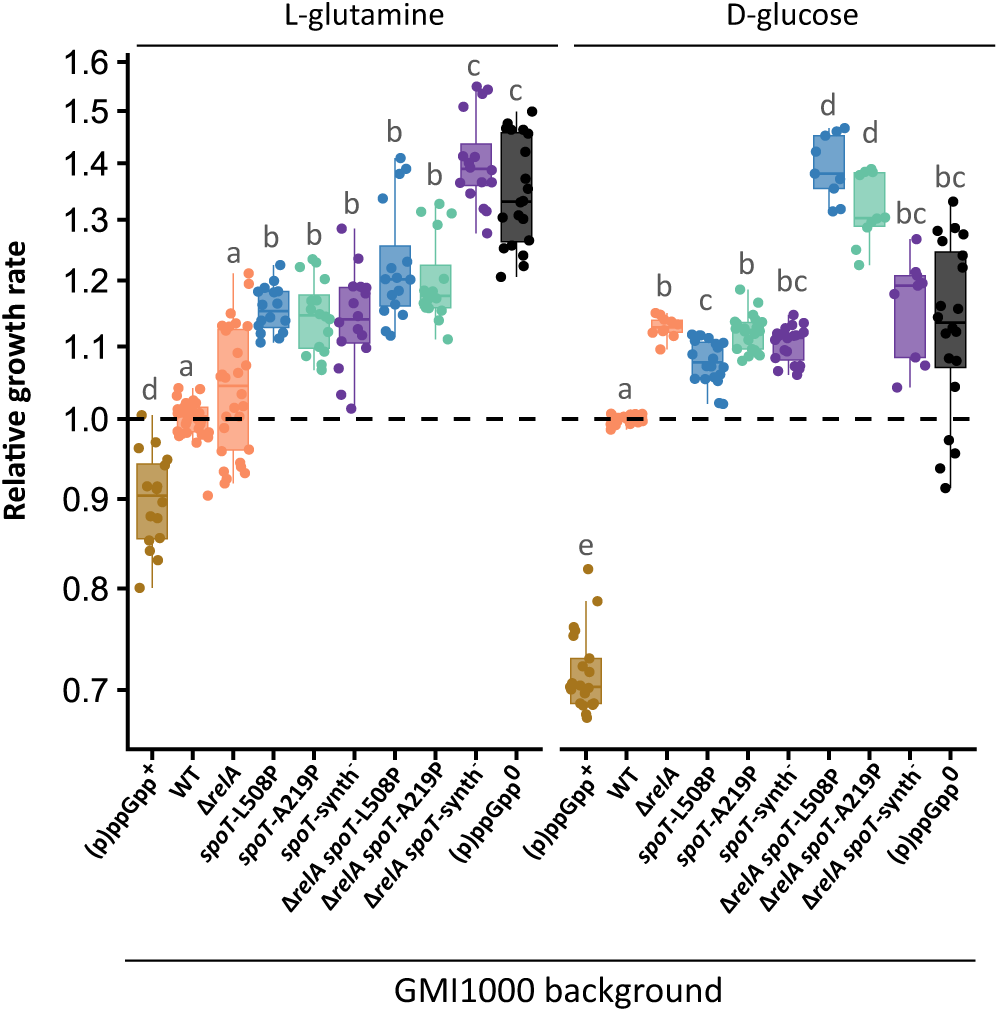
Relative growth rates of mutants affected for the synthesis of (p)ppGpp in synthetic minimal medium containing L-glutamine or D-glucose as sole carbon sources. Growth rates were measured in synthetic medium containing 10 mM L-glutamine or 56 mM D-glucose as sole carbon source. All values were normalized to the mean of the wild-type strain for each independent replicate (n=3-6), with each replicate consisting of 1-4 cultures per strain. Statistical analyses were performed using a pairwise Wilcoxon test. Different letters indicate significantly different conditions (*p* < 0.05). Raw data are available in Supplementary Table S1E. **ALT Text Fig. 4**: Two-panel figure showing boxplots with individual data points for the growth rates of strains affected in (p)ppGpp synthesis, relative to the wild-type strain, obtained from cultures grown in a synthetic medium with either glutamine or glucose as sole carbon sources. Statistical differences between groups are indicated. All *spoT* mutants grew faster whereas the (p)ppGpp⁺ strain grew more slowly than the wild-type strain.

**Table 1:** Carbon and nitrogen compounds better utilized by the GMI1000 *spoT*-A219P and GMI1000 *spoT*-L508P mutants.

| Substrate | Source | Xylem sap<sup>§</sup> | GMI1000<sup>#</sup> | *spoT*-A219P<sup>#</sup> | *spoT*-L508P<sup>#</sup> |
| --- | --- | --- | --- | --- | --- |
| L-Glutamine | C source | + | 22021 ± 494 | 24626 ± 494* (1.1) | 25010 ± 1597 (1.1) |
| D-Trehalose | C source | + | 18039 ± 1340 | 22008 ± 982* (1.2) | 21016 ± 2179 (1.2) |
| Citric Acid | C source | + | 12477 ± 1482 | 18910 ± 1151* (1.5) | 20789 ± 2050* (1.7) |
| D-Saccharic Acid | C source | + | 7627 ± 2867 | 15205 ± 3893 (2) | 17604 ± 1497* (2.3) |
| L-Histidine | C source | + | 7076 ± 2692 | 18974 ± 3642* (2.7) | 17720 ± 426* (2.5) |
| D-Galactose | C source | + | 6660 ± 1029 | 9025 ± 4984 (1.4) | 13132 ± 1459* (2) |
| Glycerol | C source | - | 6359 ± 937 | 12715 ± 3980 (2) | 13581 ± 2091* (2.1) |
| L-Alanine | C source | + | 6128 ± 1774 | 20164 ± 5285* (3.3) | 16012 ± 935* (2.6) |
| D-Gluconic Acid | C source | + | 6117 ± 2240 | 11867 ± 2846 (1.9) | 14199 ± 1245* (2.3) |
| b-Hydroxy-Butyric Acid | C source | + | 6094 ± 910 | 17723 ± 3968* (2.9) | 10588 ± 4498 (1.7) |
| L-Lactic Acid | C source | + | 5856 ± 1630 | 15853 ± 7508 (2.7) | 13423 ± 836* (2.3) |
| Sucrose | C source | + | 5472 ± 753 | 9490 ± 2954 (1.7) | 8695 ± 1226* (1.6) |
| D-Sorbitol | C source | - | 4869 ± 56 | 9695 ± 4897 (2) | 12891 ± 1688* (2.6) |
| D-Fructose | C source | + | 4596 ± 869 | 7328 ± 711* (1.6) | 11848 ± 2212* (2.6) |
| L-Serine | C source | + | 4290 ± 249 | 8507 ± 3152 (2) | 13540 ± 3720* (3.2) |
| L-Threonine | C source | + | 3763 ± 391 | 5982 ± 412* (1.6) | 9131 ± 711* (2.4) |
| L-Proline | C source | + | 3002 ± 303 | 15531 ± 9788 (5.2) | 5911 ± 475* (2) |
| Propionic Acid | C source | - | 2753 ± 745 | 5184 ± 2966 (1.9) | 9706 ± 1138* (3.5) |
| a-Keto-Butyric Acid | C source | - | 2050 ± 428 | 8002 ± 4520 (3.9) | 4061 ± 877* (2) |
| L-Histidine | N source | + | 17216 ± 1620 | 22993 ± 2639* (1.3) | 20344 ± 774 (1.2) |
| Xanthine | N source | - | 11348 ± 2274 | 16037 ± 238 (1.4) | 16465 ± 1683* (1.5) |
| Sodium Nitrite | N source | - | 7913 ± 895 | 12304 ± 1378* (1.6) | 12708 ± 119* (1.6) |
| Adenine | N source | - | 5546 ± 1304 | 11025 ± 2387* (2) | 8297 ± 1829 (1.5) |
| Met-Ala | N source | - | 5205 ± 1007 | 8831 ± 1913 (1.7) | 9704 ± 1441* (1.9) |

<sup>§</sup> Substrates detected in tomato xylem sap according to Baroukh 2022 or Lowe-Power 2018.
<sup>#</sup> Average area under the curve ± standard deviation obtained from Biolog microarrays, measuring substrate catabolism through tetrazolium dye reduction over time. Average increase ratios are shown in brackets.
\* Statistically different from GMI1000 ( $p < 0.05$ , $t$ -test). Raw data are available in Supplementary Table S1D.

### (p)ppGpp levels are likely to be reduced in the GMI1000 *spoT*-A219P and *spoT*-L508P mutants

*In vitro* phenotyping of the two *spoT* mutants showed an increased exponential growth rate in synthetic minimal media containing L-glutamine (Fig. 4). Because steady-state exponential growth rates in minimal synthetic media have been repeatedly demonstrated to be linearly and inversely correlated with basal (p)ppGpp levels in *E. coli* (Imholz et al. 2020), we hypothesized that the SpoT mutations that appeared in *R. pseudosolanacearum* changed basal (p)ppGpp levels. We therefore sought to determine how these SpoT variants affect intracellular (p)ppGpp concentrations under unstressed conditions. For comparison, we used the wild-type GMI1000 strain, a (p)ppGpp^+^ strain synthesizing elevated (p)ppGpp levels (through *relA* overexpression), a (p)ppGpp^0^ strain unable to synthesize (p)ppGpp (Δ*relA* Δ*spoT)*, and a *spoT*-synth^-^ mutant with abolished SpoT synthetase activity (*spoT*-E333Q) (Harinarayanan et al. 2008; Chau et al. 2021). Relative basal levels of ppGpp were analyzed in these strains by ion chromatography coupled to high-resolution mass spectrometry (IC-ESI-HRMS) during exponential growth in modified low-phosphate M9 medium (Patacq et al. 2018) containing 1% glucose (56 mM). In this medium, the *spoT*-A219P, *spoT*-L508P, *spoT*-synth^-^ and (p)ppGpp^0^ strains grew slightly but significantly faster than the wild-type strain whereas the (p)ppGpp^+^ strain grew much more slowly (Fig. S2AB). Relative quantification of ppGpp revealed that the (p)ppGpp^+^ strain harbored 36-fold higher ppGpp levels than the wild-type strain, while no ppGpp was detected in the (p)ppGpp^0^ strain.

However, ppGpp levels measured in the wild-type strain and in the *spoT-*A219P, *spoT-*L508P and *spoT-*synth^-^ mutants were close to the detection limit (near background noise) and could not be statistically distinguished (Fig. S2C).

As a complementary approach to assess basal (p)ppGpp levels in *Ralstonia* strains, we measured the exponential growth rate of a set of mutants altered in (p)ppGpp synthesis. In addition to the (p)ppGpp^+^, the (p)ppGpp^0^ and the *spoT*-synth^-^ (E333Q) strains described above, we introduced the three *spoT* mutations (A219P, L508P, and E333Q) into a Δ*relA* background, in which SpoT was the sole (p)ppGpp synthetase. Consequently, the Δ*relA spoT-*synth*^-^* mutant is theoretically equivalent to a (p)ppGpp^0^ strain, as it lacks any functional (p)ppGpp synthetase. To ensure comparability, all strains including the wild-type strain, carried the same gentamicin resistance gene. The genomes of all *spoT* mutants were sequenced, and the strains were complemented with the wild-type *spoT* allele to check for the absence of compensatory mutations. Exponential growth rates were first measured in rich medium, in which most amino acids and growth factors are available. In these conditions, most mutant strains exhibited growth rates similar to, or slightly slower than, that of the wild-type strain. Only the (p)ppGpp^+^ and (p)ppGpp^0^ mutants showed significantly reduced growth rates compared to the wild-type strain (Fig. S3). Growth rates were then measured in a synthetic minimal medium containing either 10 mM L-glutamine, the main carbon source used by *R. pseudosolanacearum* in xylem sap (Baroukh et al. 2022), or 56 mM D-glucose, as sole carbon sources (Fig. 4). As expected, the (p)ppGpp^+^ strain grew more slowly than the wild-type strain in both minimal media. More surprisingly, the (p)ppGpp^0^ strains (Δ*relA* Δ*spoT* and Δ*relA spoT-*synth^-^) grew faster than the wild-type strain and even exhibited the fastest generation times in minimal medium with L-glutamine as the sole carbon source. This contrasts with observations in *E. coli* and *Bacillus subtilis*, where (p)ppGpp^0^ strains fail to grow in minimal medium due to amino-acid auxotrophy (Potrykus et al. 2011; Kriel et al. 2014). In *Pseudomonas putida* and *Cupriavidus necator* (formerly *Ralstonia eutropha*), (p)ppGpp^0^ strains also grow more slowly than the wild-type while retaining prototrophy (Juengert et al. 2017; Vogeleer and Létisse 2022). The ability of the *R. pseudosolanacearum* (p)ppGpp^0^ strains to grow in an amino acid-free minimal synthetic medium (with D-glucose as sole carbon source) indicates that they retain the capacity to synthesise all amino acids.

In L-glutamine minimal medium, the *R. pseudosolanacearum spoT*-A219P, *spoT*-L508P, and *spoT*-synth^-^ mutants displayed intermediate growth rates between those of the wild-type and (p)ppGpp^0^ strains (Fig. 4). Deletion of *relA* did not significantly increase the growth rate, indicating that SpoT is the main (p)ppGpp synthetase in this medium. Consistently, the *spoT*-A219P and *spoT*-L508P mutants exhibited similar growth rates in both the wild-type and Δ*relA* backgrounds in L-glutamine minimal medium, but significantly lower growth rates than the Δ*relA spoT*-synth^-^ strain. We confirmed that these growth phenotypes were due to the *spoT* mutations, as complementation with the wild-type *spoT* allele restored the wild-type growth phenotype (Fig. S4).

In D-glucose minimal medium, the *spoT*-A219P, *spoT*-L508P, and *spoT*-synth^-^ mutants also exhibited higher growth rates than the wild-type strain, comparable to those of the (p)ppGpp^0^ strains (Fig. 4). In this medium, however, RelA appears to contribute to basal (p)ppGpp synthesis, as deletion of *relA* significantly increased the growth rate. Moreover, combining either the *spoT*-A219P or *spoT*-L508P mutation with the *relA* deletion resulted in an additive effect on growth, with double mutants growing even faster than the (p)ppGpp^0^ strains on this carbon source.

Altogether, growth rate measurements in minimal medium suggest that the *spoT*-A219P and *spoT*-L508P mutations reduce basal (p)ppGpp levels without completely abolishing SpoT synthetase activity.

### Inactivation of the SpoT synthetase activity enhances the fitness of bacteria *in planta*

To determine whether inactivation of the SpoT (p)ppGpp synthetase activity is adaptive *in planta*, we assessed the fitness of the *spoT*-synth^-^ mutants in tomato xylem and during symbiosis with *M. pudica*, either in competition with the wild-type strain or in single strain inoculation assays. In both experimental setups, the *spoT*-synth⁻ mutants behaved similarly to the *spoT*-A219P and *spoT*-L508P mutants. In co-inoculations, it exhibited enhanced *in planta* fitness compared with the wild-type strain in both plant environments (Fig. 2AC). In single strain inoculations, the number of bacteria per gram of tomato stem did not differ significantly between the wild-type and the *spoT*-synth⁻ mutant five days post-inoculation (DPI). In contrast, in *M. pudica* nodules, the *spoT*-synth⁻ mutant showed significantly higher proliferation than the wild-type strain. We also evaluated the *in planta* adaptiveness of the (p)ppGpp^0^ (Δ*relA* Δ*spoT*) strain in tomato xylem. In co-inoculation assays with the wild-type strain, this mutant displayed enhanced competitive fitness, as did the other *spoT* mutants. However, it exhibited slightly reduced xylem colonisation in single strain inoculations at 5 DPI (see Fig. 2AB) and reduced virulence on tomato plants following root inoculation or stem injection (Fig. 3). The phenotype of a (p)ppGpp^0^ strain could not be assessed during the interaction with *M. pudica*, as we were unable to generate the Δ*relA* Δ*spoT* deletions in the nodulating background (GMI1000 pRalta *hrpG*). Conversely, the strain that constitutively expressed *relA* and produced higher basal levels of (p)ppGpp exhibited significantly reduced fitness when competing with the wild-type strain in both tomato xylem and *M. pudica* nodules (Fig. 2AC). This strain also exhibited reduced virulence on tomato and induced poorly infected nodules on *M. pudica* in single strain inoculations (Fig. 2D, Fig. 3).

Together, these results indicate that the complete suppression of (p)ppGpp synthesis or the inactivation of SpoT synthetase activity improves bacterial fitness in both plant xylem and legume nodules. This supports the idea that the adaptive advantage conferred by the *spoT*-A219P and *spoT*-L508P mutations may be due to lower basal (p)ppGpp levels. Although analyses of the (p)ppGpp^0^ strain show that (p)ppGpp is not strictly required for xylem colonization, our data further revealed that optimal pathogenicity of the *R. pseudosolanacearum* GMI1000 strain relies on finely tuned (p)ppGpp levels, neither absent nor too high, and that elevated basal levels of (p)ppGpp is detrimental to legume symbiosis in this system.

### The enhanced growth rate of the *spoT* mutants may explain their increased fitness in plants

As basal (p)ppGpp levels have recently been shown to coordinate metabolic homeostasis and acid resistance in *E. coli* (Liu et al. 2026), and given that xylem sap and nodule symbiosomes are likely to be acidic (Jia and Davies 2007; Pierre et al. 2013), we also evaluated the effect of pH on the growth of the mutants. Growth was assessed in a minimal medium adjusted to pH 4.5, 5.0, 5.5 or 6.0 and compared with growth at pH 6.5, the optimal pH for *R. pseudosolanacearum* GMI1000 (Peyraud et al. 2016). In these experiments, D-glucose was used as the carbon source because the pH increased rapidly when L-glutamine was used, unlike D-glucose, which maintained a low pH (not shown; Liu et al. 2022). No growth was observed at pH 4.5 or 5.0 (Fig. S5), and growth was very limited at pH 5.5 for all strains. At pH 6.0, the *spoT* mutants grew faster than the wild-type strain, in a manner similar to that observed at pH 6.5. These results suggest that acidic pH neither limits nor enhances the fitness advantage of the *spoT* mutants.

To test whether improved growth rates could be sufficient to explain the enhanced fitness of the *spoT* mutants in xylem, we used the measured growth rates in minimal medium containing L-glutamine to simulate their expected fitness, based on the estimated number of bacterial generations occurring in xylem vessels (Guidot et al. 2014). The competitive indexes predicted from these growth differences fit relatively well the values measured *in planta* (Fig. S6), suggesting that improved utilization of L-glutamine may be one of the most important factors responsible for the increased fitness of the mutants in xylem.

Overall, these results indicate that mutations in *spoT* have been selected primarily for their enhanced metabolic and growth abilities, which probably contributed to their improved fitness in both plant environments.

## Discussion

Evolution experiments have repeatedly demonstrated that bacteria often adapt through mutations in global regulators (Cooper et al. 2003; Maharjan et al. 2006; Philippe et al. 2007; Wang et al. 2010). Such mutations, which rewire regulatory networks, generally have large fitness effects and therefore play a critical role during the earliest stages of adaptation (Barrick et al. 2009; Capela et al. 2017; Lenski 2017). Because these mutations are pleiotropic, they may confer benefits across multiple environments. This is for example the case of mutations in the RNA polymerase subunits *rpoB* and *rpoC*, which enable adaptation to both elevated temperature and antibiotic exposure in *E. coli* (Rodríguez-Verdugo et al. 2013). In this study, we demonstrated that two independent mutations in the *spoT* gene of *R. pseudosolanacearum* confer a fitness advantage in two distinct plant environments, the xylem of plants, both susceptible and tolerant to *R. pseudosolanacearum*, and the root nodules of legumes. The mechanisms underlying this enhanced fitness remain unclear. Rather than being a specific adaptation to plant responses, these mutations may enable bacteria to better cope with the physiological conditions prevailing in these environments. In particular, plants may provide specific nutrients that bacteria have to metabolize efficiently. Biolog phenotyping revealed that the A219P and L508P mutations in *spoT* enhanced the ability of strain GMI1000 to utilize several carbon and nitrogen compounds. For example, sugars and amino acids such as glutamine, proline and glucose, which are present at millimolar concentrations in tomato xylem sap (Baroukh et al. 2022), represent potential carbon sources that were more efficiently metabolized by the mutants. Consistent with this observation, we confirmed that the *spoT* mutants grow faster than the wild-type strain when L-glutamine is provided as the sole carbon source. L-glutamine is not only one of the most abundant compounds in tomato xylem sap but is also commonly found in the xylem sap of other plant species (Hocking 1980; Andersen and Brodbeck 1989; Gerlin et al. 2021), and appears to be the preferred substrate assimilated by *R. pseudosolanacearum* GMI1000 in this compartment (Baroukh et al. 2022). Moreover, our simulation analyses indicate that the increased growth rate of the *spoT* mutants relative to the wild-type strain in this minimal medium containing L-glutamine could be sufficient to account for their enhanced competitive fitness in xylem. Although the metabolites present in *Mimosa* nodules remain largely unknown, previous studies have shown that the host plant supplies symbiotic bacteria with several amino acids. In particular, branched-chain amino acids, aspartate and glutamate have been reported to be essential for some rhizobium-legume interactions (Lodwig et al. 2003; Prell et al. 2009; Flores-Tinoco et al. 2020; Wu et al. 2026). Interestingly, in previous work we identified mutations affecting the EfpR regulatory pathway that were highly adaptive in both plant xylem and legume nodules. These mutations also markedly increased the ability of the bacteria to utilize a broad range of carbon and nitrogen substrates that were otherwise poorly metabolized by the wild-type strain (Perrier et al. 2016; Capela et al. 2017). This shared phenotype between *efpR* and *spoT* mutants further support the idea that improving nutrient utilization is a key factor in the adaptation of bacteria to these plant environments.

The role of (p)ppGpp has long been associated with the stringent response, characterized by a rapid induction of high levels of the molecule. However, it is now clear that (p)ppGpp production in bacteria is gradual and varies with stress intensity (Spira and Ospino 2020; Steinchen et al. 2020; Engelhardt et al. 2025). For example, Zhu et al. (2024) showed that a moderate increase in basal level of (p)ppGpp in *E. coli* maintains exponential growth at a moderate rate while significantly enhancing stress tolerance. In addition, low basal levels of (p)ppGpp are also known to play an important regulatory role as a secondary messenger in the physiology of bacteria, modulating balanced growth rates and maintaining the homeostasis of important cellular components and macromolecules, as reviewed by (Fernández-Coll and Cashel 2020). Minor adjustments to these basal levels can have profound phenotypic consequences, affecting not only growth (Zhu and Dai 2019) but also motility and cell population organization (Ababneh and Herman 2015) or acid stress resistance (Liu et al. 2026). Our results suggest that the A219P and L508P adaptive mutations in *spoT* reduce basal (p)ppGpp level, as evidenced by the increased growth rate of the mutants under unstressed conditions and similar *in vitro* and *in planta* phenotypes to those observed in a *spoT*-synth^-^ mutant impaired in the SpoT synthetase activity. Although this reduction could not be demonstrated through direct measurement of ppGpp levels using mass spectrometry, it cannot be ruled out that these experiments only measured free ppGpp, rather than ppGpp molecules that were effectively bound to cellular partners, such as proteins or ribosomes. Moreover, in a Δ*relA* background the *spoT*-A219P and *spoT*-L508P mutants are not equivalent to the *spoT*-synth^-^ mutant suggesting that these mutants are likely to be only partially affected in synthetase activity. This may imply that even a slight reduction in (p)ppGpp levels could confer significant adaptive advantages in the tested plant environments. Fine-tuned basal concentrations of (p)ppGpp were also shown to be critical for virulence in animal pathogens where (p)ppGpp^0^ mutants generally exhibit attenuated virulence (Weiss and Stallings 2013; Spira et al. 2014; Kundra et al. 2020). These basal levels are essential for both acute and chronic infection in *Mycobacterium tuberculosis* as well as for macrophage transmission of *Legionella pneumophila* for example (Dalebroux et al. 2010; Weiss and Stallings 2013). In plant-associated bacteria, while basal (p)ppGpp levels have not been specifically studied, the synthesis of (p)ppGpp is required for both pathogenic and mutualistic interactions. Indeed (p)ppGpp^0^ strains are typically impaired in their interaction with plants. For example, in the plant pathogens *Xanthomonas campestris*, *Xanthomonas citri*, *Pseudomonas syringae* or *Dickeya dantatii*, (p)ppGpp regulates essential virulence-associated genes including genes encoding Type III secretion systems, EPS biosynthesis or cell wall degrading enzymes and (p)ppGpp^0^ mutants of these strains display reduced virulence (Ancona et al. 2015; Chatnaparat et al. 2015a; Chatnaparat et al. 2015b; Cui et al. 2019; Zhang et al. 2019; Bai et al. 2021; Shi et al. 2025). We have shown here that the virulence of a (p)ppGpp^0^ derivative strain of *R. pseudosolanacearum* GMI1000 is reduced. Similarly, (p)ppGpp^0^ mutants of the legume symbionts *Sinorhizobium meliloti* and *Rhizobium etli* are either unable to infect the root tissues of their host plants or are altered in their capacity to fix nitrogen in nodules (Calderón-Flores et al. 2005; Moris et al. 2005; Wippel and Long 2019). Interestingly, in *R. pseudosolanacearum*, the complete absence of (p)ppGpp is less detrimental than previously reported for other bacteria. In *E. coli* and *B. subtilis*, (p)ppGpp^0^ strains exhibit amino acid auxotrophy and severe defects in biosynthetic pathways (Potrykus et al. 2011; Kriel et al. 2014). In other species, such as *P. putida* and *C. necator*, (p)ppGpp^0^ strains remain prototrophic but show slower growth in minimal medium (Juengert et al. 2017; Vogeleer and Létisse 2022). This contrasts sharply with the *R. pseudosolanacerum* (p)ppGpp^0^ strain, which not only grew in media lacking essential growth factors (vitamins and amino-acids) and even displayed the highest growth rate of all the strains in this study when grown in minimal medium containing L-glutamine as carbon source. Remarkably, this strain was also more fit than the wild-type strain in competition for xylem colonization, which was unexpected given that (p)ppGpp^0^ mutants in other species are generally impaired in host interactions, as discussed above. These findings confirm that, although (p)ppGpp-regulated functions are broadly conserved across bacteria, the regulation and targets of (p)ppGpp have been slightly reshaped in certain species during evolution, resulting in species-specific differences.

The two *spoT* mutations affect amino acid residues that are strictly conserved across the *Ralstonia solanacearum* species complex and widely conserved in other bacteria (Atkinson et al. 2011). This high level of conservation suggests the presence of selective pressures that have prevented such mutations from arising or being maintained in natural populations, raising the possibility of trade-offs between the fitness benefits observed in the xylem and potential costs during other stages of the bacterial life cycle. Notably, the two *spoT* mutants showed no impairment in virulence following root infection, a critical step in the *R. pseudosolanacearum* life cycle that was bypassed during our experimental evolution experiment, in which bacteria were propagated by direct reinjection into plant stems (Guidot et al. 2014). In the literature, (p)ppGpp levels have been proposed to regulate global resource allocation by balancing cellular investment between ribosome synthesis, which promotes rapid growth, and the production of metabolic and stress-related proteins, which generally reduces growth rate while enhancing stress resistance (Ferenci and Spira 2007; Zhu and Dai 2019; Fernández-Coll and Cashel 2020; Zhu et al. 2024). In environments where nutrients are continuously available, such as xylem sap (Baroukh et al. 2022), strong investment in stress-response pathways or specific biosynthetic functions may be less critical. Consistent with this idea, a universal trade-off between maximal growth rate and the time required to adapt to a new environment has been described (Basan et al. 2020). In our evolution experiment, where bacteria were maintained under relatively stable conditions in xylem sap, the selective pressure to rapidly adapt to fluctuating environments was likely relaxed, potentially favoring specialization to this niche. In contrast, under natural conditions *R. pseudosolanacearum* must cope with drastic lifestyle transitions between the xylem and nutrient-poor environments such as soil or environmental water, which constitute major reservoirs of *Ralstonia* and where bacteria can experience prolonged starvation (Álvarez et al. 2008). Transition to these extreme conditions may require increased (p)ppGpp synthesis (Patacq et al. 2020; Zhu et al. 2024). In such contexts, the *spoT* mutations identified here may no longer be advantageous, which could explain why these amino acid residues remain highly conserved in natural populations.

Noteworthy, mutations in *spoT* frequently arise in adaptive laboratory evolution (ALE) experiments. The ALEdb 1.0 database reports 70 distinct SNPs in *spoT* from eleven ALE experiments involving four strains and 528 samples (Phaneuf et al. 2019). However, these data are biased towards an over-representation of evolutionary studies of *E. coli* in *in vitro* culture conditions. For instance, mutations in *spoT* were detected in 8 different lineages over 12 of the well-known Lenski’s evolution experiment that serially propagated *E. coli* in glucose-limited conditions for more than 80,000 generations (Cooper et al. 2003; Philippe et al. 2007) as well as in *E. coli* evolved under phosphate limitation (Wang et al. 2010). Hence, the *spoT* gene is considered a mutational hotspot and ranks as the fourth most frequently targeted gene by adaptive mutations in *E. coli* evolution studies (Tenaillon et al. 2016; Wang et al. 2018). However, mutations in *spoT* were never reported for bacteria evolving in contact with an eukaryotic host. Moreover, the physiological impact of most of the *spoT* mutations from previous ALE experiments were not determined.

In conclusion, the recurrent selection of different *spoT* alleles in evolution experiments suggests that fine-tuning of intrinsic concentrations of (p)ppGpp is an efficient strategy for bacteria to optimize adaptation to specific environments. Depending on nutrient availability, physiological demands and stress exposure, either a slight increase or decrease in (p)ppGpp levels may confer a selective advantage. Our study demonstrated that a reduction of basal (p)ppGpp levels in *R. pseudosolanacearum* enhanced bacterial fitness in plant-associated environments, including the xylem sap of both susceptible and tolerant hosts and the nodules of legumes. Given the central and pleiotropic role of (p)ppGpp in coordinating transcription, translation, metabolism and growth, pinpointing the exact molecular mechanisms underlying these adaptations remains challenging. One possible explanation for the benefits of the *spoT* mutants is their ability to utilize more rapidly key carbon and nitrogen sources. Alternatively, the SpoT-mediated basal regulation of (p)ppGpp levels may coordinate metabolism with stress resistance as recently described for acid stress resistance in *E. coli* (Liu et al. 2026). Nevertheless, our results support the idea that small shifts in basal (p)ppGpp concentrations can produce large phenotypic effects and enable bacteria to rapidly adjust to specific ecological niches. More broadly, this work emphasizes the importance of quantitative modulation of global regulatory systems in shaping bacterial adaptation to complex host-associated environments.

## Materials and methods

### Bacterial strains and growth conditions

All strains used in this study are listed into the supplementary material Table S4A. *R. pseudosolanacearum* strains were cultivated at 28°C in either rich medium (per liter: 10 g Bactopeptone; 1 g casamino acid; 1 g yeast extract) or synthetic minimal medium (per liter: 3.4 g KH2PO4; 0.5 g (NH4)2SO4; 0.05 g MgSO4,7H2O) supplemented with a carbon source (either L-glutamine 10mM or 56 mM D-glucose for growth assays or 2% (v/v) glycerol for bacterial transformation) and trace elements (per liter: 1.5 mg Na2EDTA,2H2O; 4.5 mg ZnSO4,7H2O; 0.3 mg CoCl2,6H2O; 1 mg MnCl2,4H2O; 1 mg H3BO3; 0.4 mg Na2MoO4,2H2O; 3 mg FeSO4,7H2O; 0.3 mg CuSO4,5H2O) and adjusted to pH=6.5 (unless otherwise stated) using 1 M KOH. *Escherichia coli* strains were cultivated on Luria-Bertani medium at 37°C. Antibiotics were used at the following concentrations: trimethoprim 100 µg.mL^-1^, spectinomycin 40 µg.mL^-1^, gentamicin 10 µg.mL^-1^, tetracycline 10 µg.mL^-1^.

### Genome sequencing of evolved clones and detection of genomic modifications

Evolved clones were grown overnight in rich medium, centrifugated at 5,000 g for 10 minutes, before the pellets being frozen and kept at -80°C. DNA extractions were performed using the Wizard® genomic DNA purification kit (Promega). DNAs of clones evolved on *M. pudica* were sequenced at the GeT-PlaGe core facility (https://get.genotoul.fr/), INRAE Toulouse using the Illumina technology and mutations were detected as previously described (Marchetti et al. 2017). DNAs of clones evolved on cabbage were sequenced in this work at Genoscope (https://jacob.cea.fr/drf/ifrancoisjacob/english/Pages/Departments/Genoscope.aspx) using the Illumina Technology and mutations were detected using the open-source *breseq* computational pipeline (Deatherage and Barrick 2014).

### Conservation of amino-acid positions across various bacterial species

The SpoT (RSc2153) protein sequence was used as a query in BLASTp searches against protein datasets from 120 *Ralstonia* genomes to identify SpoT homologs. A global multiple sequence alignment was performed using MAFFT (v7.505) on the 121 proteins to assess sequence conservation. A phylogenetic tree was inferred from the SpoT protein sequences using an automated phylogenetic pipeline to examine their evolutionary relationships using phylogeny.fr (Dereeper et al. 2008).

### Strain constructions

All PCR primers and plasmids used for this work are listed in Table S4BC. The genomes of all the *spoT* mutants constructed in the GMI1000 genetic background used in this study were resequenced using the Nanopore technology to detect potential other mutations in the genome. The ONT Nanopore reads were mapped to the reference genome of *R. pseudosolanacearum* GMI1000 using minimap2 (v2.30-r187) with ’map-ont’ preset. Alignments shorter than 10 kb were filtered out before polymorphism analysis with samtools. SNPs and small indels were detected with VarScan mpileup2cns (v2.4.6), while structural variants were detected with CuteSV (v2.0.3) using the parameters recommended by the authors for ONT data (Jiang et al. 2020).

The A219P and L508P point mutations in *spoT* were reconstructed into the ancestral strain using the MuGENT technique (Multiplex Genome Editing by Natural Transformation) (Dalia et al. 2014) as detailed in Capela et al. (2017). Briefly, competent cells were obtained after a 48 h culture in minimal medium with 2% (v/v) glycerol and incubated 48 h onto rich medium with 3 µg of a 6 kb fragment harboring the mutation and 300 ng of an integrative linearised plasmid carrying a gentamicine-resistance selection cassette and a gene encoding a fluorescent protein (*Sca*I-linearized pRCG-*PpsbA-GFP* or pRCG-*PpsbA-mCherry* plasmids adapted from Monteiro et al. 2012, that integrate in the intergenic region downstream the *glmS* gene). The 6 kb fragments were amplified from evolved clones using the Phusion DNA polymerase. Transformants were selected on gentamicine and screened for the presence of the SNP mutation by PCR. The 3’-end of one primer was designed to align with the mutated or wild-type nucleotide, while a mismatch was introduced at the penultimate nucleotide to increase stringency. Integration was verified through Sanger or Nanopore sequencing.

Deletions of *relA* (*RSc1576*) or *spoT* were performed as described in Huang and Wilks (2017). Around 800 bp fragments were amplified upstream and downstream of the target sequence, and cloned together into the suicide plasmid pEX18Tc (Choi and Schweizer 2005) using the *HindIII*/*XbaI* and *XbaI*/*BamHI* restriction sites. Plasmids were introduced into *Ralstonia* strains by natural transformation. After chromosomal integration of the pEX18Tc suicide plasmids carrying the *sacB* gene, bacteria were cultivated on rich medium with 5% (m/v) sucrose, and deletion mutants were screened using regular PCR amplification. To construct the (p)ppGpp^0^ strain, the *relA* gene was deleted prior to the deletion of *spoT*. Once the *relA* deletion was constructed in GMI1000, a 7 kb fragment around the deletion was amplified using the Phusion polymerase, and used to delete the gene in the *spoT-*A219P and *spoT*-L508P mutants using the MuGENT procedure described above.

Overexpression of *relA* was achieved by integrating a copy of the *relA* gene into the intergenic region downstream of the *glmS* gene under the control of the constitutive *PpsbA* promoter (Brixey et al. 1997; Puigvert et al. 2019). To achieve this, the *GFP* gene from the pRCG-*PpsbA*-GFP plasmid (Monteiro et al. 2012) was replaced with the *relA* gene through double digestion of the plasmid with *Spe*I and *Acc65I* restriction enzymes and ligation with overhang PCR digested with the same restriction enzymes. The *relA* constructions were inserted into *R. pseudosolanacearum* GMI1000 strain by natural transformation.

The *spoT*-synth^-^ mutant was constructed by substituting glutamic acid at position 333 with glutamine (mutation E333Q). This mutation targets a conserved amino acid essential for the SpoT synthetase catalytic site, previously described in *E. coli* (mutation E319Q) (Harinarayanan et al. 2008) and *Salmonella enterica* (Chau et al. 2021). The mutated gene *spoT*-E333Q was cloned into the pEX18Tc plasmid using two overhang PrimeStar PCR fragments (Takara Bio Inc.). The two fragments of approximately 800 bp were amplified downstream and upstream of the mutation site, each containing a 20 bp overlap with the other fragment, with the desired mismatch mutation in the center of the overlap region. Each fragment also harbored a 20 bp overhang overlap with one of the *SacI*/*SalI*-digested ends of the pEX18Tc plasmid. The two fragments and the linearized plasmid were then assembled through the complementary sequences using the In-Fusion cloning kit (Takara Bio Inc.). Integration was first realized into the GMI1000 Δ*relA* strain, followed by plasmid excision using the *sacB*-mediated selection method. Then, a 6 kb fragment around the *spoT*-E333Q mutation was amplified from GMI1000 Δ*relA spoT*-E333Q, and used to introduce this mutation in the GMI1000 strain by the MuGent method as detailed above for the introduction of point mutations.

To complement the *spoT* mutants, the wild-type *spoT* allele was amplified by PCR using a high-fidelity PrimeSTAR DNA polymerase (Takara Bio Inc.) and primers extended with 15 bp complementary with *SacI*/*SalI*-digested ends of the pEX18Tc plasmid. The PCR product and the linearized plasmid were then assembled using the In-Fusion cloning kit (Takara Bio Inc.) and transformed into *E. coli* competents cells. The plasmid was then integrated into the native *spoT* locus by natural transformation of *R. pseudosolanacearum* strains and subsequently excised using the *sacB*-mediated selection method. Complemented strains were identified by PCR screening and verified by nanopore sequencing of the *spoT* locus.

### Plant assays

Tomato (*Solanum lycopersicum* var. Marmande) and cabbage (*Brassica oleracea* var. Bartolo) plants were cultivated in a greenhouse for 4–5 weeks before being transferred to a growth chamber at 27–28 °C with 80% humidity and a 12-hour light/dark cycle. *Mimosa pudica* plants were grown in test tubes following the protocol described in (Doin de Moura et al. 2023) in a chamber at 28°C and a 16-hour light/dark cycle.

For competition assays in the xylem of tomato and cabbage, plants were co-inoculated with pairs of strains carrying the *GFP* and *mCherry* fluorochrome encoding genes, respectively. To prepare the 1:1 inoculum, strains were separately cultivated overnight in rich medium, then diluted in sterile water to a final concentration of 10⁶ CFU.mL^-1^ and mixed in equal volumes. Five plants per mix were inoculated by direct stem injection of 15 µL. Bacteria were sampled after 5 days by cutting the plant stem, surface sterilizing with 70% ethanol, and submerging in 3 mL of sterile water to allow bacteria to diffuse out of the stem. For competition assays on *M. pudica*, bacteria grown on solid agar plates were resuspended in sterile water, adjusted to 2 × 10⁷ CFU.mL^-1^, and mixed. 100 µL of bacterial mix was inoculated into each growth tube containing two *M. pudica* plantlets. Nodules were harvested from 20 plants after 21 days, pooled, surface sterilized in 2.4° bleach for 15 minutes, and crushed.

The bacterial suspensions were plated on rich medium agar using an easySpiral automatic plater (Interscience, France). Colonies expressing mCherry and GFP fluorochromes were counted after 48 hours of incubation at 28°C using a ZEISS V16 fluorescent stereo-zoom microscope. The competitive indexes (CIs) were calculated as the final ratio divided by the inoculum ratio (Morel et al. 2018). When one of the two strains was undetectable, ratios were calculated supposing 1 colony on the counted plate.

The predicted competitive indexes of the mutants in the xylem were calculated using the formula CI_predicted_=2^n*(µmax_*spoT*-µmax_*wt*)/µmax_*wt*^, n being the number of bacterial generations measured 5 days post-inoculation, µmax_*spoT* and µmax_WT the calculated growth rates in minimal medium containing L-glutamine as the sole carbon source (see below). In tomato Marmande, the number of generations after one passage (5 days post-inoculation) was estimated at 13 in our previous work (Guidot et al. 2014).

For the measurement of bacterial infectivity in *M. pudica* nodules, 10^6^ bacteria were inoculated onto 5-day-old seedlings. Ten plants per strain were used. Nodules were harvested 21 days post inoculation, surface-sterilized, crushed, diluted and plated as described above. For the measurement of infectivity in tomato plants, 10^4^ bacteria were injected directly into the stem of five plants. After five days, plants were harvested, and bacteria were recovered from around 1 g of fresh stem material, as described above and plated.

For nodule infected area quantification, *M. pudica* nodules were harvested at 21 days post inoculation (DPI) and cut into 55 µm sections with a vibrating blade microtome (VT1000 S, Leica, Wetzlar, Germany). Quantification of infection areas on nodule sections was performed as described in Marchetti et al. (2014) using the Image-Pro Plus software and based on the HIS method (Media Cybernetics, Rockville, MD, USA). Results of quantifications of infected areas were obtained from 2 to 3 independent experiments.

For pathogenicity assays on tomatoes, two methods were used to inoculate plants. For inoculation by soil drenching, overnight cultures in rich medium were diluted to 5 × 10⁷ CFU.mL^-1^ in 500 mL of sterile water and evenly distributed over the soil of a tray containing 16 plants. For stem inoculation, 10 µL of the bacterial culture diluted to 10⁶ CFU.mL^-1^ in sterile water were injected into the stem just above the cotyledons of each of 16 plants in a tray. Symptoms were monitored daily and scored on a scale of 0 to 4, following a scale described in Morel et al. 2018. For survival curve analysis, a plant was recorded as dead when symptoms reached a score of 3 (75% of the leaves wilted) or higher (complete wilting). Survival curves and hazard ratios (HR) were calculated using the Mantel-Haenszel method in the GraphPad software.

### *In vitro* phenotyping

Metabolic profiles were assessed using the Biolog phenotyping microarrays PM01, PM02, PM03. Plates were prepared following the manufacturer’s instructions (Biolog, Inc., Hayward, CA) and incubated at 28°C in the Omnilog reader for 96 hours. Analysis has been realized using the *opm* package in the R studio software to extract the area under the curves (AUC).

For growth monitoring, strains were pre-cultivated overnight in the assay medium and diluted to OD600nm of 0.05. 200 µL of each culture was then dispensed into five wells of a 96-well plate, and OD600nm was then measured every 6.6 minutes in an Omega Plate Reader (FLUOstar, BMG Labtech), shaking at 700 rpm for 50-100 hours. Growth rates were then assessed using a linear regression on the growth curve after transformation by natural logarithm in the R Studio software.

### ppGpp quantification

Analysis of ppGpp was performed following the protocol described in Patacq et al. (2018). Strains were grown in a modified low-phosphate M9 medium (per liter: 3.92 g Na₂HPO₄·12H₂O, 0.606 g KH₂PO₄, 0.51 g NaCl, 2.04 g NH₄Cl, 0.098 g MgSO₄, 4.38 mg CaCl₂) supplemented with 0.1 g·L⁻¹ thiamine hydrochloride, trace elements (per liter: 1.5 mg Na₂EDTA·2H₂O, 4.5 mg ZnSO₄·7H₂O, 0.3 mg CoCl₂·6H₂O, 1 mg MnCl₂·4H₂O, 1 mg H₃BO₃, 0.4 mg Na₂MoO₄·2H₂O, 3 mg FeSO₄·7H₂O, 0.3 mg CuSO₄·5H₂O), and 56 mM D-glucose. The pH of the cultures was manually adjusted to 6.5 using 1 M NaOH at two time points: 1.5 h after inoculation (OD₆₀₀ = 0.05–0.15) and during exponential phase (OD₆₀₀ ≃ 0.5). Cultures were sampled during exponential growth and metabolic activity was rapidly quenched using a cold solution (methanol/acetonitrile/H₂O, 4:4:2). After centrifugation at 10,000 × g for 10 min, supernatants were collected and solvents were removed using a SpeedVac for 4–6 h. The remaining aqueous phase was then lyophilized and stored at −70 °C. Prior to analysis, samples were reconstituted in ultrapure water and analyzed using ion chromatography (IC; Thermo Scientific Dionex ICS-5000+, Dionex, Sunnyvale, CA, USA) coupled to a Q Exactive Orbitrap mass spectrometer (Thermo Fisher Scientific, Waltham, MA, USA) equipped with a heated electrospray ionization (HESI) source. Separation was achieved using a KOH gradient as follows: 7 mM (0–1 min), linearly increased to 15 mM (1–9.5 min), held at 15 mM (9.5–20 min), then ramped to 45 mM (20–30 min), to 70 mM (30–33 min), and to 100 mM (33.1–42 min), followed by re-equilibration at 7 mM (42.5–50 min) (Jaboulay et al. 2025). The total run time was 50 min, and 10 µL were injected. Mass spectra were acquired in negative ion mode using full-scan acquisition at a resolution of 70,000 (m/z 200), with the following source parameters: capillary temperature 300 °C, heater temperature 450 °C, sheath gas flow 40 AU, auxiliary gas flow 20 AU, S-lens RF level 60%, and spray voltage −3.5 kV. Metabolites were identified based on exact m/z values with a 5 ppm mass tolerance, using Xcalibur software (Thermo Fisher Scientific) and processed with El-Maven (version 0.12.0) (Agrawal et al. 2019). Three independent biological replicates were analyzed.

## Supplementary Materials

**Table S1.** Raw data generated in this study used to produce the main and supplementary figures.

**Table S2.** List of mutations detected in clones evolved in Cabbage, lineage C.

**Table S3.** List of mutations detected in clones evolved in *M. pudica*, lineage D.

**Table S4.** Lists of strains, plasmids and primers used in this study.

**Figure S1.** Conservation of the A219 and L508 amino-acid position across the *Ralstonia solanacearum* species complex.

**Figure S2.** Bacterial growth in low-phosphate M9 minimal medium and relative quantification of ppGpp

**Figure S3.** Growth rates of mutants affected for the synthesis of (p)ppGpp in rich medium

**Figure S4.** Growth rates of the complemented mutants with the wild-type *spoT* allele in minimal synthetic medium with L-glutamine as sole carbon sources.

**Figure S5.** Growth of *R. pseudosolanacearum* GMI1000 mutants harboring different (p)ppGpp levels at different pH

**Figure S6.** Comparison of predicted and observed competitive indexes (CIs) of the *R. pseudosolanacearum* GMI1000 *spoT* and (p)ppGpp+ mutants in tomato xylem

## Acknowledgements

N.B. was supported by a fellowship from the French Ministère de l’Enseignement Supérieur, de la Recherche et de l’Innovation (MESRI). This study was supported by the French National Research Agency (ANR-22-CE20-0014), the “Laboratoires d’Excellence (LABEX)” TULIP (ANR-10-LABX-41), and the “École Universitaire de Recherche (EUR)” TULIP-GS (ANR-18-EURE-0019).

## Author contributions

N.B., A.P., A.G. and D.C. conceptualized research. N.B. M.T., L.L, F.L, P.V and D.C. performed experiments. N.B. analyzed and curated all data. A.G. and D.C. provided supervision. N.B., A.G. and D.C. wrote the original draft. All authors reviewed and approved the manuscript.

## Data availability

All raw data are provided in Table S1.

## Notes

### Competing Interest Statement

The authors have declared no competing interest.

## References

1. Ababneh QO, Herman JK. 2015. RelA inhibits *Bacillus subtilis* motility and chaining. J Bacteriol. 197:128–137.

2. Agrawal S, Kumar S, Sehgal R, George S, Gupta R, Poddar S, Jha A, Pathak S. 2019. El-MAVEN: A fast, robust, and user-friendly mass spectrometry data processing engine for metabolomics. In : High-Throughput Metabolomics. Vol. 1978. Methods in Molecular Biology. New York, NY: Springer New York. p. 301–321. doi:10.1007/978-1-4939-9236-2_19

3. Álvarez B, López MM, Biosca EG. 2008. Survival strategies and pathogenicity of *Ralstonia solanacearum* phylotype II subjected to prolonged starvation in environmental water microcosms. Microbiology 154:3590–3598.

4. Amadou C, Pascal G, Mangenot S, Glew M, Bontemps C, Capela D, Carrère S, Cruveiller S, Dossat C, Lajus A, et al. 2008. Genome sequence of the β-rhizobium *Cupriavidus taiwanensis* and comparative genomics of rhizobia. Genome Res. 18:1472–1483.

5. Ancona V, Lee JH, Chatnaparat T, Oh J, Hong J-I, Zhao Y. 2015. The bacterial alarmone (p)ppGpp activates the type III secretion system in *Erwinia amylovora*. J Bacteriol. 197:1433–1443.

6. Andersen PC, Brodbeck BV. 1989. Diurnal and temporal changes in the chemical profile of xylem exudate from *Vitis rotundifolia*. Physiologia Plantarum 75:63–70.

7. Atkinson GC, Tenson T, Hauryliuk V. 2011. The RelA/SpoT Homolog (RSH) Superfamily: distribution and functional evolution of ppGpp synthetases and hydrolases across the tree of life. PLoS ONE 6:e23479.

8. Bai K, Yan H, Chen X, Lyu Q, Jiang N, Li J, Luo L. 2021. The role of RelA and SpoT on ppGpp production, stress response, growth regulation, and pathogenicity in *Xanthomonas campestris* pv. *campestris*. Microbiol Spectr. 9:e02057–21.

9. Baroukh C, Zemouri M, Genin S. 2022. Trophic preferences of the pathogen *Ralstonia solanacearum* and consequences on its growth in xylem sap. MicrobiologyOpen 11. doi:10.1002/mbo3.1240

10. Barrick JE, Yu DS, Yoon SH, Jeong H, Oh TK, Schneider D, Lenski RE, Kim JF. 2009. Genome evolution and adaptation in a long-term experiment with *Escherichia coli*. Nature 461:1243–1247.

11. Basan M, Honda T, Christodoulou D, Hörl M, Chang Y-F, Leoncini E, Mukherjee A, Okano H, Taylor BR, Silverman JM, et al. 2020. A universal trade-off between growth and lag in fluctuating environments. Nature 584:470–474.

12. Bergsma-Vlami M, Van De Bilt JLJ, Tjou-Tam-Sin NNA, Westenberg M, Meekes ETM, Teunissen HAS, Van Vaerenbergh J. 2018. Phylogenetic assignment of *Ralstonia pseudosolanacearum* (*Ralstonia solanacearum* Phylotype I) isolated from *Rosa* spp. Plant Disease 102:2258–2267.

13. Briones G, Iñón De Iannino N, Roset M, Vigliocco A, Paulo PS, Ugalde RA. 2001. *Brucella abortus* cyclic β-1,2-glucan mutants have reduced virulence in mice and are defective in intracellular replication in HeLa cells. Infect Immun. 69:4528–4535.

14. Brixey PJ, Guda C, Daniell H. 1997. The chloroplast *psbA* promoter is more efficient in *Escherichia coli* than the T7 promoter for hyperexpression of a foreign protein. Biotechnology Letters 19:395–400.

15. Calderón-Flores A, Du Pont G, Huerta-Saquero A, Merchant-Larios H, Servín-González L, Durán S. 2005. The stringent response is required for amino acid and nitrate utilization, Nod Factor regulation, nodulation, and nitrogen fixation in *Rhizobium etli*. J Bacteriol. 187:5075–5083.

16. Campbell GRO, Reuhs BL, Walker GC. 2002. Chronic intracellular infection of alfalfa nodules by *Sinorhizobium meliloti* requires correct lipopolysaccharide core. Proc Natl Acad Sci U S A. 99:3938–3943.

17. Capela D, Marchetti M, Clérissi C, Perrier A, Guetta D, Gris C, Valls M, Jauneau A, Cruveiller S, Rocha EPC, et al. 2017. Recruitment of a lineage-specific virulence regulatory pathway promotes intracellular infection by a plant pathogen experimentally evolved into a legume symbiont. Mol Biol Evol. 34:2503–2521.

18. Chatnaparat T, Li Z, Korban SS, Zhao Y. 2015a. The stringent response mediated by (p)ppGpp is required for virulence of *Pseudomonas syringae* pv. *tomato* and its survival on tomato. Mol Plant Microbe Interact. 28:776–789.

19. Chatnaparat T, Li Z, Korban SS, Zhao Y. 2015b. The bacterial alarmone (p)ppGpp is required for virulence and controls cell size and survival of *Pseudomonas syringae* on plants. Environ Microbiol. 17:4253–4270.

20. Chau NYE, Pérez-Morales D, Elhenawy W, Bustamante VH, Zhang YE, Coombes BK. 2021. (p)ppGpp-dependent regulation of the nucleotide hydrolase PpnN confers complement resistance in *Salmonella enterica* serovar *Typhimurium*. Infect Immun. 89:e00639–20.

21. Choi K-H, Schweizer HP. 2005. An improved method for rapid generation of unmarked *Pseudomonas aeruginosa* deletion mutants. BMC Microbiol. 5:30.

22. Cooper TF, Rozen DE, Lenski RE. 2003. Parallel changes in gene expression after 20,000 generations of evolution in *Escherichia coli*. Proc Natl Acad Sci U S A. 100:1072–1077.

23. Cui Z, Yang C-H, Kharadi RR, Yuan X, Sundin GW, Triplett LR, Wang J, Zeng Q. 2019. Cell-length heterogeneity: a population-level solution to growth/virulence trade-offs in the plant pathogen *Dickeya dadantii*. bioRxiv. doi:10.1101/577296

24. Dalebroux ZD, Yagi BF, Sahr T, Buchrieser C, Swanson MS. 2010. Distinct roles of ppGpp and DksA in Legionella pneumophila differentiation. Mol Microbiol. 76(1):200–219.

25. Dalia AB, McDonough E, Camilli A. 2014. Multiplex genome editing by natural transformation. Proc Natl Acad Sci U S A. 111:8937–8942.

26. Deatherage DE, Barrick JE. 2014. Identification of mutations in laboratory-evolved microbes from next-generation sequencing data using breseq. Engineering and analyzing multicellular systems. Vol. 1151. Methods in Molecular Biology. New York, NY: Springer New York. p. 165–188. doi:10.1007/978-1-4939-0554-6_12

27. Dereeper A, Guignon V, Blanc G, Audic S, Buffet S, Chevenet F, Dufayard J-F, Guindon S, Lefort V, Lescot M, et al. 2008. Phylogeny.fr: robust phylogenetic analysis for the non-specialist. Nucleic Acids Res. 36:W465–W469.

28. Doin de Moura GG, Mouffok S, Gaudu N, Cazalé A-C, Milhes M, Bulach T, Valière S, Roche D, Ferdy J-B, Masson-Boivin C, et al. 2023. A selective bottleneck during host entry drives the evolution of new legume symbionts. Mol Biol Evol. 40(5):msad116.

29. Doin de Moura GG, Remigi P, Masson-Boivin C, Capela D. 2020. Experimental evolution of legume symbionts: what have we learnt? Genes 11:339.

30. Drew GC, Stevens EJ, King KC. 2021. Microbial evolution and transitions along the parasite–mutualist continuum. Nat Rev Microbiol. 19:623–638.

31. Engelhardt F, Turnbull K, Gür M, Müsken M, Preusse M, Häussler S, Roghanian M. 2025. (p)ppGpp imposes graded transcriptional changes to impair motility and promote antibiotic tolerance in biofilms. npj Biofilms Microbiomes 11:148.

32. Ferenci T, Spira B. 2007. Variation in stress responses within a bacterial species and the indirect costs of stress resistance. Ann N Y Acad Sci. 1113:105–113.

33. Fernández-Coll L, Cashel M. 2020. Possible Roles for Basal levels of (p)ppGpp: growth efficiency vs. surviving stress. Front Microbiol. 11:592718.

34. Flores-Tinoco CE, Tschan F, Fuhrer T, Margot C, Sauer U, Christen M, Christen B. 2020. Co-catabolism of arginine and succinate drives symbiotic nitrogen fixation. Mol Syst Biol. 16:e9419.

35. Genin S, Denny TP. 2012. Pathogenomics of the *Ralstonia solanacearum* Species Complex. Annu Rev Phytopathol. 50:67–89.

36. Gentry DR, Cashel M. 1996. Mutational analysis of the *Escherichia coli spoT* gene identifies distinct but overlapping regions involved in ppGpp synthesis and degradation. Mol Microbiol. 19:1373–1384.

37. Gerlin L, Escourrou A, Cassan C, Maviane Macia F, Peeters N, Genin S, Baroukh C. 2021. Unravelling physiological signatures of tomato bacterial wilt and xylem metabolites exploited by *Ralstonia solanacearum*. Environ Microbiol. 23:5962–5978.

38. Gopalan-Nair R, Jardinaud M-F, Legrand L, Landry D, Barlet X, Lopez-Roques C, Vandecasteele C, Bouchez O, Genin S, Guidot A. 2021. Convergent rewiring of the virulence regulatory network promotes adaptation of *Ralstonia solanacearum* on resistant tomato. Mol Biol Evol. 38:1792–1808.

39. Gopalan-Nair R, Jardinaud M-F, Legrand L, Lopez-Roques C, Bouchez O, Genin S, Guidot A. 2023. Transcriptomic profiling reveals host-specific evolutionary pathways promoting enhanced fitness in the plant pathogen *Ralstonia pseudosolanacearum*. Microbial Genomics 9. doi:10.1099/mgen.0.001142

40. Guidot A, Jiang W, Ferdy J-B, Thébaud C, Barberis P, Gouzy J, Genin S. 2014. Multihost experimental evolution of the pathogen *Ralstonia solanacearum* unveils genes involved in adaptation to plants. Mol Biol Evol. 31:2913–2928.

41. Harinarayanan R, Murphy H, Cashel M. 2008. Synthetic growth phenotypes of *Escherichia coli* lacking ppGpp and transketolase A (*tktA*) are due to ppGpp-mediated transcriptional regulation of *tktB*. Mol Microbiol. 69:882–894.

42. Hauryliuk V, Atkinson GC, Murakami KS, Tenson T, Gerdes K. 2015. Recent functional insights into the role of (p)ppGpp in bacterial physiology. Nat Rev Microbiol. 13:298–309.

43. Hocking PJ. 1980. The composition of phloem exudate and xylem sap from tree tobacco (*Nicotiana glauca* Grah.). Ann Bot. 45:633–643.

44. Huang W, Wilks A. 2017. A rapid seamless method for gene knockout in *Pseudomonas aeruginosa*. BMC Microbiol. 17:199.

45. Imholz NCE, Noga MJ, Van Den Broek NJF, Bokinsky G. 2020. Calibrating the bacterial growth rate speedometer: a re-evaluation of the relationship between basal ppGpp, growth, and RNA synthesis in *Escherichia coli*. Front Microbiol. 11:574872.

46. Irving SE, Choudhury NR, Corrigan RM. 2021. The stringent response and physiological roles of (pp)pGpp in bacteria. Nat Rev Microbiol. 19:256–271.

47. Jaboulay C, Dugelay C, Maio JPS, Beillevaire K, Vogeleer P, Vinella D, Létisse F, Hallez R, Wang B, Terradot L, et al. 2025. Cross-regulation of (p)ppGpp and c-di-GMP pathways controls a cell-cycle transition. bioRxiv. doi:10.1101/2025.08.22.671821

48. Jia W, Davies WJ. 2007. Modification of leaf apoplastic pH in relation to stomatal sensitivity to root-sourced abscisic acid signals. Plant Physiol. 143:68–77.

49. Jiang T, Liu Yongzhuang, Jiang Y, Li J, Gao Y, Cui Z, Liu Yadong, Liu B, Wang Y. 2020. Long-read-based human genomic structural variation detection with cuteSV. Genome Biol. 21:189.

50. Juengert JR, Borisova M, Mayer C, Wolz C, Brigham CJ, Sinskey AJ, Jendrossek D. 2017. Absence of ppGpp leads to increased mobilization of intermediately accumulated Poly(3-Hydroxybutyrate) in *Ralstonia eutropha* H16. Appl Environ Microbiol. 83:e00755–17.

51. Kriel A, Brinsmade SR, Tse JL, Tehranchi AK, Bittner AN, Sonenshein AL, Wang JD. 2014. GTP dysregulation in *Bacillus subtilis* cells lacking (p)ppGpp results in phenotypic amino acid auxotrophy and failure to adapt to nutrient downshift and regulate biosynthesis genes. J Bacteriol. 196:189–201.

52. Kundra S, Colomer-Winter C, Lemos JA. 2020. Survival of the fittest: The relationship of (p)ppGpp with bacterial virulence. Front Microbiol. 11:601417.

53. Lenski RE. 2017. Experimental evolution and the dynamics of adaptation and genome evolution in microbial populations. ISME J. 11:2181–2194.

54. LeVier K, Phillips RW, Grippe VK, Roop RM, Ii, Walker GC. 2000. Similar requirements of a plant symbiont and a mammalian pathogen for prolonged intracellular survival. Science 287:2492–2493.

55. Liu Y, Schicketanz ML, Zhai X, Deng L, Gerdes K, Zhang YE. 2026. Basal ppGpp signalling by SpoT integrates metabolism with acid resistance. bioRxiv. doi:10.64898/2026.01.26.700336.

56. Liu Y, Tan X, Pan Y, Yu J, Du Y, Liu X, Ding W. 2022. Mutation in phcA enhanced the adaptation of *Ralstonia solanacearum* to long-term acid stress. Front Microbiol. 13:829719.

57. Lodwig EM, Hosie AHF, Bourdès A, Findlay K, Allaway D, Karunakaran R, Downie JA, Poole PS. 2003. Amino-acid cycling drives nitrogen fixation in the legume–Rhizobium symbiosis. Nature 422:722–726.

58. Lomovatskaya LA, Romanenko AS. 2020. Secretion systems of bacterial phytopathogens and mutualists. Appl Biochem Microbiol. 56:115–129.

59. Lowe-Power TM, Hendrich CG, Von Roepenack-Lahaye E, Li B, Wu D, Mitra R, Dalsing BL, Ricca P, Naidoo J, Cook D, et al. 2018. Metabolomics of tomato xylem sap during bacterial wilt reveals *Ralstonia solanacearum* produces abundant putrescine, a metabolite that accelerates wilt disease. Environ Microbiol. 20:1330–1349.

60. Maddamsetti R, Hatcher PJ, Green AG, Williams BL, Marks DS, Lenski RE. 2017. Core genes evolve rapidly in the long-term evolution experiment with *Escherichia coli*. Genome Biol Evol. 9:1072–1083.

61. Maharjan R, Seeto S, Notley-McRobb L, Ferenci T. 2006. Clonal Adaptive radiation in a constant environment. Science 313:514–517.

62. Mansfield J, Genin S, Magori S, Citovsky V, Sriariyanum M, Ronald P, Dow M, Verdier V, Beer SV, Machado MA, et al. 2012. Top 10 plant pathogenic bacteria in molecular plant pathology. Mol Plant Pathol. 13:614–629.

63. Marchetti M, Capela D, Glew MD, Cruveiller S, Chane-Woon-Ming B, Béatrice Chane-Woon-Ming, Gris C, Timmers T, Poinsot V, Véréna Poinsot, et al. 2010. Experimental evolution of a plant pathogen into a legume symbiont. PLoS Biol. 8(1):e1000280.

64. Marchetti M, Clerissi C, Yousfi Y, Gris C, Bouchez O, Rocha EPC, Cruveiller S, Jauneau A, et al. 2017. Experimental evolution of rhizobia may lead to either extra- or intracellular symbiotic adaptation depending on the selection regime. Mol Ecol. 26:1818–1831.

65. Marchetti M, Jauneau A, Capela D, Remigi P, Gris C, Batut J, Masson-Boivin C. 2014. Shaping bacterial symbiosis with legumes by experimental evolution. Mol Plant Microbe Interact. 27:956–964.

66. Mechold U, Potrykus K, Murphy H, Murakami KS, Cashel M. 2013. Differential regulation by ppGpp versus pppGpp in *Escherichia coli*. Nucleic Acids Res. 41:6175–6189.

67. Melnyk RA, Hossain SS, Haney CH. 2019. Convergent gain and loss of genomic islands drive lifestyle changes in plant-associated *Pseudomonas*. ISME J. 13:1575–1588.

68. Monteiro F, Solé M, Van Dijk I, Valls M. 2012. A chromosomal insertion toolbox for promoter probing, mutant complementation, and pathogenicity studies in *Ralstonia solanacearum*. Mol Plant Microbe Interact. 25:557–568.

69. Morel A, Peeters N, Vailleau F, Barberis P, Jiang G, Berthomé R, Guidot A. 2018. Plant pathogenicity phenotyping of *Ralstonia solanacearum* strains. Host-Pathogen Interactions. Vol. 1734. Methods in Molecular Biology. New York, NY: Springer New York. p. 223–239. doi:10.1007/978-1-4939-7604-1_18

70. Moris M, Braeken K, Schoeters E, Verreth C, Beullens S, Vanderleyden J, Michiels J. 2005. Effective symbiosis between *Rhizobium etli* and *Phaseolus vulgaris* requires the alarmone ppGpp. J Bacteriol. 187:5460–5469.

71. Morris CE, Moury B. 2019. Revisiting the concept of host range of plant pathogens. Annu Rev Phytopathol. 57:63–90.

72. Patacq C, Chaudet N, Létisse F. 2018. Absolute quantification of ppGpp and pppGpp by double-spike isotope dilution ion chromatography-high-resolution mass spectrometry. Anal Chem. 90:10715–10723.

73. Patacq C, Chaudet N, Létisse F. 2020. Crucial role of ppGpp in the resilience of *Escherichia coli* to growth disruption. mSphere 5:e01132–20.

74. Perrier A, Peyraud R, Rengel D, Barlet X, Lucasson E, Gouzy J, Peeters N, Genin S, Guidot A. 2016. Enhanced in planta fitness through adaptive mutations in EfpR, a dual regulator of virulence and metabolic functions in the plant pathogen *Ralstonia solanacearum*. PLoS Pathog. 12:e1006044.

75. Peyraud R, Cottret L, Marmiesse L, Gouzy J, Genin S. 2016. A resource allocation trade-off between virulence and proliferation drives metabolic versatility in the plant pathogen *Ralstonia solanacearum*. PLoS Pathog. 12:e1005939.

76. Phaneuf PV, Gosting D, Palsson BO, Feist AM. 2019. ALEdb 1.0: a database of mutations from adaptive laboratory evolution experimentation. Nucleic Acids Res. 47:D1164–D1171.

77. Philippe N, Crozat E, Lenski RE, Schneider D. 2007. Evolution of global regulatory networks during a long- term experiment with *Escherichia coli*. BioEssays 29:846–860.

78. Pierre O, Engler G, Hopkins J, Brau F, Boncompagni E, Hérouart D. 2013. Peribacteroid space acidification: a marker of mature bacteroid functioning in *Medicago truncatula* nodules: Acidification of peribacteroid space in root nodule. Plant Cell Environ. doi: 10.1111/pce.12116

79. Potrykus K, Cashel M. 2008. (p)ppGpp: still magical? Annu Rev Microbiol. 62:35–51.

80. Potrykus K, Murphy H, Philippe N, Cashel M. 2011. ppGpp is the major source of growth rate control in *E. coli*: ppGpp and growth rate control. Environ Microbiol. 13:563–575.

81. Prell J, White JP, Bourdès A, Bunnewell S, Bongaerts RJ, et al. Legumes regulate Rhizobium bacteroid development and persistence by the supply of branched-chain amino acids. 2009. Proc Natl Acad Sci U S A. 106:12477–12482. doi: 10.1073/pnas.0903653106.

82. Puigvert M, Solé M, López-Garcia B, Coll NS, Beattie KD, Davis RA, Elofsson M, Valls M. 2019. Type III secretion inhibitors for the management of bacterial plant diseases. Mol Plant Pathol. 20:20–32.

83. Richardson EJ, Bacigalupe R, Harrison EM, Weinert LA, Lycett S, Vrieling M, Robb K, Hoskisson PA, Holden MTG, Feil EJ, et al. 2018. Gene exchange drives the ecological success of a multi-host bacterial pathogen. Nat Ecol Evol. 2:1468–1478.

84. Rodríguez-Verdugo A, Gaut BS, Tenaillon O. 2013. Evolution of *Escherichia coli* rifampicin resistance in an antibiotic-free environment during thermal stress. BMC Evol Biol. 13:50.

85. Ronneau S, Hallez R. 2019. Make and break the alarmone: regulation of (p)ppGpp synthetase/hydrolase enzymes in bacteria. FEMS Microbiol Rev. 43:389–400.

86. Sachs JL, Skophammer RG, Regus JU. 2011. Evolutionary transitions in bacterial symbiosis. Proc Natl Acad Sci U S A. 108:10800–10807.

87. Sanchez-Vazquez P, Dewey CN, Kitten N, Ross W, Gourse RL. 2019. Genome-wide effects on *Escherichia coli* transcription from ppGpp binding to its two sites on RNA polymerase. Proc Natl Acad Sci U S A. 116:8310–8319.

88. Shi Z, Liang Z, Yang Q, Zhang L-H, Wang Q. 2025. Alarmone ppGpp modulates bacterial motility, zeamine production, and virulence of *Dickeya oryzae* through the regulation of and cooperation with the putrescine signaling mechanism. mSphere 10:e00682–24.

89. Sinha AK, Winther KS, Roghanian M, Gerdes K. 2019. Fatty acid starvation activates RelA by depleting lysine precursor pyruvate. Mol Microbiol. 112:1339–1349.

90. Smith NT, Boukherissa A, Antaya K, Howe GW, Mergaert P, Rodríguez De La Vega RC, Shykoff JA, Alunni B, diCenzo GC. 2025. Taxonomic distribution of SbmA/BacA and BacA-like antimicrobial peptide transporters suggests independent recruitment and convergent evolution in host–microbe interactions. Microbial Genomics 11. doi:10.1099/mgen.0.001380.

91. Soto MJ, Sanjuán J, Olivares J. 2006. Rhizobia and plant-pathogenic bacteria: common infection weapons. Microbiology (Reading) 152, 3167–3174.

92. Spira B, Ferreira GM, De Almeida LG. 2014. relA enhances the adherence of enteropathogenic *Escherichia coli*. PLoS ONE 9:e91703.

93. Spira B, Ospino K. 2020. Diversity in *E. coli* (p)ppGpp Levels and its consequences. Front Microbiol. 11:1759.

94. Steinchen W, Zegarra V, Bange G. 2020. (p)ppGpp: Magic modulators of bacterial physiology and metabolism. Front Microbiol. 11:2072.

95. Tamman H, Ernits K, Roghanian M, Ainelo A, Julius C, Perrier A, Talavera A, Ainelo H, Dugauquier R, Zedek S, et al. 2023. Structure of SpoT reveals evolutionary tuning of catalysis via conformational constraint. Nat Chem Biol. 19(3):334–345.

96. Tenaillon O, Barrick JE, Ribeck N, Deatherage DE, Blanchard JL, Dasgupta A, Wu GC, Wielgoss S, Cruveiller S, Médigue C, et al. 2016. Tempo and mode of genome evolution in a 50,000-generation experiment. Nature 536:165–170.

97. Teulet A, Camuel A, Perret X, Giraud E. 2022. The versatile roles of type III secretion systems in Rhizobium-Legume symbioses. Annu. Rev. Microbiol. 76:45–65.

98. Vogeleer P, Létisse F. 2022. Dynamic metabolic response to (p)ppGpp accumulation in *Pseudomonas putida*. Front Microbiol. 13:872749.

99. Wang L, Spira B, Zhou Z, Feng L, Maharjan RP, Li X, Li F, McKenzie C, Reeves PR, Ferenci T. 2010. Divergence involving global regulatory gene mutations in an *Escherichia coli* population evolving under phosphate limitation. Gen Biol Evol. 2:478–487.

100. Wang X, Zorraquino V, Kim M, Tsoukalas A, Tagkopoulos I. 2018. Predicting the evolution of *Escherichia coli* by a data-driven approach. Nat Commun. 9:3562.

101. Weiss LA, Stallings CL. 2013. Essential roles for *Mycobacterium tuberculosis* Rel beyond the production of (p)ppGpp. J Bacteriol. 195:5629–5638.

102. Wicker E, Grassart L, Coranson-Beaudu R, Mian D, Prior P. 2009. Epidemiological evidence for the emergence of a new pathogenic variant of *Ralstonia solanacearum* in Martinique (French West Indies). Plant Pathol. 58:853–861.

103. Wippel K, Long SR. 2019. Symbiotic performance of *Sinorhizobium meliloti* lacking ppGpp depends on the *Medicago* host species. Mol Plant Microbe Interact. 32:717–728.

104. Wu J, Zhang X, Fan Z, Huang Y, Cao Y, Ren J, Yang L, Tian J, Yu Y, Kong Z. 2026. Symbiosome membrane-localized cationic amino acid transporters support symbiotic nitrogen fixation in *Medicago truncatula*. Plant Commun. 7:101636.

105. Zhang Y, Teper D, Xu J, Wang N. 2019. Stringent response regulators (p)ppGpp and DksA positively regulate virulence and host adaptation of *Xanthomonas citri*. Molecular Plant Pathol. 20:1550–1565.

106. Zhu M, Dai X. 2019. Growth suppression by altered (p)ppGpp levels results from non-optimal resource allocation in *Escherichia coli*. Nucleic Acids Res. 47:4684–4693.

107. Zhu M, Mu H, Dai X. 2024. Integrated control of bacterial growth and stress response by (p)ppGpp in *Escherichia coli*: A seesaw fashion. iScience 27:108818.

108. Zhu M, Pan Y, Dai X. 2019. (p)ppGpp: the magic governor of bacterial growth economy. Curr Genet. 65:1121–1125.

