## Supplementary figures for "SpoT-mediated reduction of (p)ppGpp levels promotes *Ralstonia pseudosolanacearum* adaptation to both plant xylem and legume nodules"

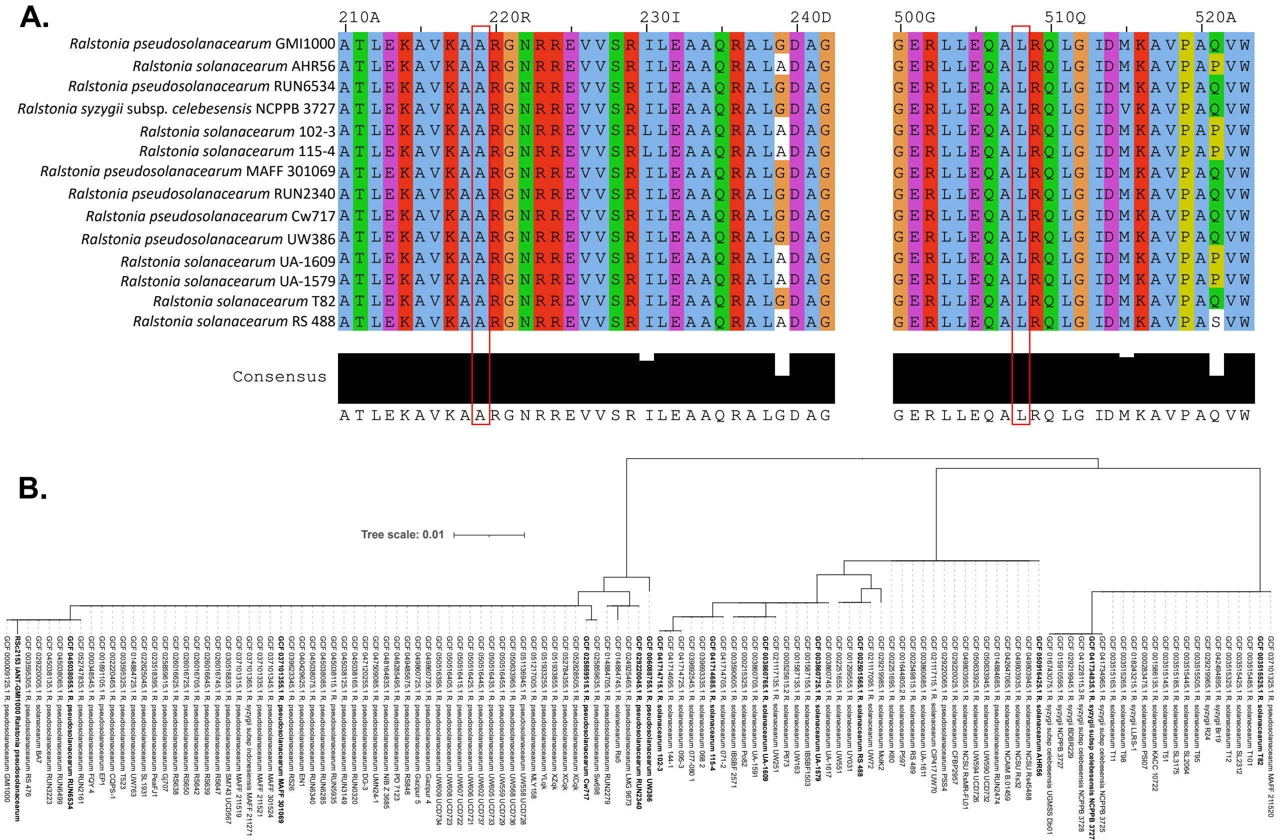


**Figure S1. Conservation of the A219 and L508 amino-acid position across the *Ralstonia solanacearum* species complex. A. Multiple alignment** of the SpoT proteins from 14 representative *Ralstonia solanacearum* strains out of the 121 analyzed. The A219 and L508 positions were conserved in all 121 SpoT proteins analyzed. **B. Phylogenetic tree** showing the 121 SpoT proteins, the sequences used for the alignment in A are highlighted in bold.

**Alt text**: Panel A shows a multiple sequence alignment of SpoT proteins from 14 representative strains of the *Ralstonia solanacearum* species complex, centered on the A219 and L508 positions, with a consensus sequence shown below and conserved residues highlighted. Panel B shows a phylogenetic tree of the 121 analyzed SpoT proteins, with the sequences used for the alignment in panel A indicated in bold. The A219 and L508 positions are conserved across all SpoT proteins examined.

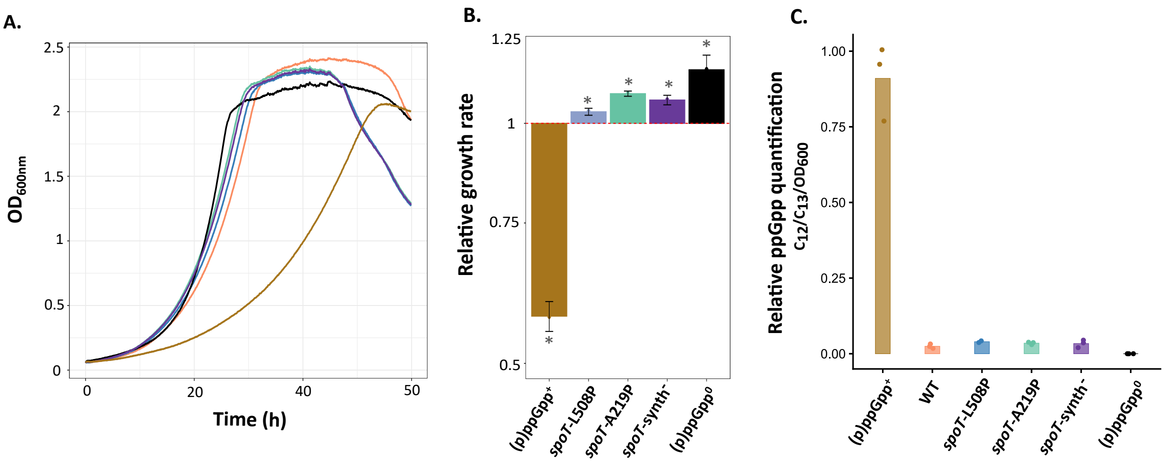


**Figure S2. Bacterial growth in low-phosphate M9 minimal medium and quantification of ppGpp**

**A. Growth curves** for one representative experiment, different colors represent different mutants as in panel **(B)** and **(C). B. Relative growth rates** in modified M9 medium containing 56 mM D-glucose. Growth rates were normalized by the growth rate of the wild-type (WT) strain for each independent replicate.* indicates growth rates significantly different to 1 ($p < 0.05$, t-test). Error bars indicate standard deviation. **C. Relative ppGpp quantification** using IC-ESI-HRMS (chromatography coupled to the electrospray ionization high-resolution mass spectrometry) on samples from cultures in exponential phase of growth. **B. C.** Data are from 2-3 independent experiments. Raw data are available in Supplementary Table S1F.

**ALT Text:** Three-panel figure showing bacterial growth and ppGpp quantification in a low-phosphate M9 medium.

The first panel is a line graph of $OD_{600}$ over time (0–50 hours) representing growth curves for (p)ppGpp$^+$, wild type, *spoT* mutants (L508P, A219P, synth$^-$), and (p)ppGpp$^0$ strains. All strains except the (p)ppGpp$^+$ strain exhibit visible enhanced growth compared to the WT. The (p)ppGpp$^0$ strain exhibits the fastest growth. The (p)ppGpp$^+$ strain has the most different growth curve. The second panel presents a bar plot showing relative growth rates and the standard deviation for the same strains normalized to the wild type (dashed line at 1). The (p)ppGpp$^+$ strain exhibits reduced growth rate, while the *spoT* mutants and the (p)ppGpp$^0$ strain exhibit increased growth. Significant differences from the WT are indicated by asterisks. Finally, the third panel displays a bar plot showing the levels of intracellular relative ppGpp (C12/C13) (normalized to $OD_{600}$), revealing high relative accumulation in the (p)ppGpp$^+$ strain, very low levels in the wild type and *spoT* mutants, and a complete absence in the (p)ppGpp$^0$ strain.

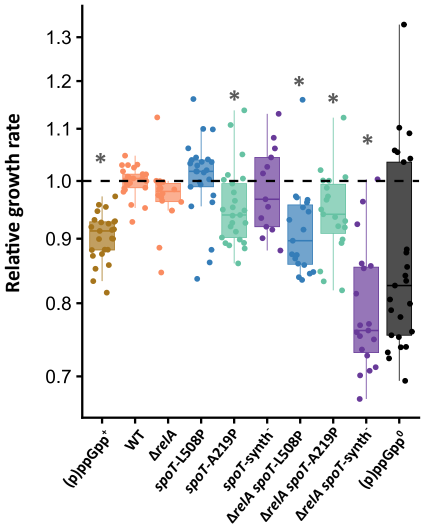


**Figure S3. Relative growth rates of mutants affected for the synthesis of (p)ppGpp in complete rich medium**

Growth rates were measured in complete rich medium. All values were normalized to the mean of the wild-type strain for each independent replicate (n=4-7), with each replicate consisting of 3-4 cultures per strain. * Significantly different from the wild-type (WT) strain (Wilcoxon test, $p < 0.05$). Raw data are available in Supplementary Table S1E.

**ALT Text**: Figure showing boxplots with individual data points for the growth rates of mutants for (p)ppGpp synthesis relative to the wild-type strain. Relative growth rate varies from 0.7 to 1.2, with most strains harboring a comparable growth rate to the WT, at the exception of the *spoT*-A219P, (p)ppGpp$^{+}$, *spoT*-L508P and Δ*relA spoT*-A219P strains that are slightly reduced in growth rate in this medium.

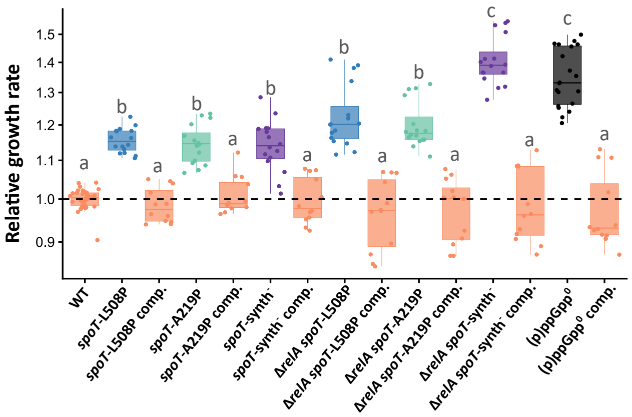


**Figure S4. Growth rates of the complemented mutants with the wild-type *spoT* allele in minimal synthetic medium with L-glutamine as sole carbon source**

Relative growth rate of complemented *spoT* strains in the GMI1000 background. All values were normalized to the mean of the wild-type strain for each independent replicate (n=3), with each replicate consisting of 4 cultures per strain. Statistical analyses were performed using a pairwise Wilcoxon test. Different letters indicate significantly different conditions ($p < 0.05$). Data for the non-complemented strains are the same as in Figure 4. Complemented strains are indicated as "comp." Raw data are available in Supplementary Table S1E.

ALT Text: Two-panel figure showing boxplots with individual data points for the growth rates of all *spoT* mutants and complemented strains relative to the wild-type strain. Statistical differences between groups are indicated. All complemented strains exhibit growth rates comparable to the wild type and thus significantly different from their parental non-complemented strain.

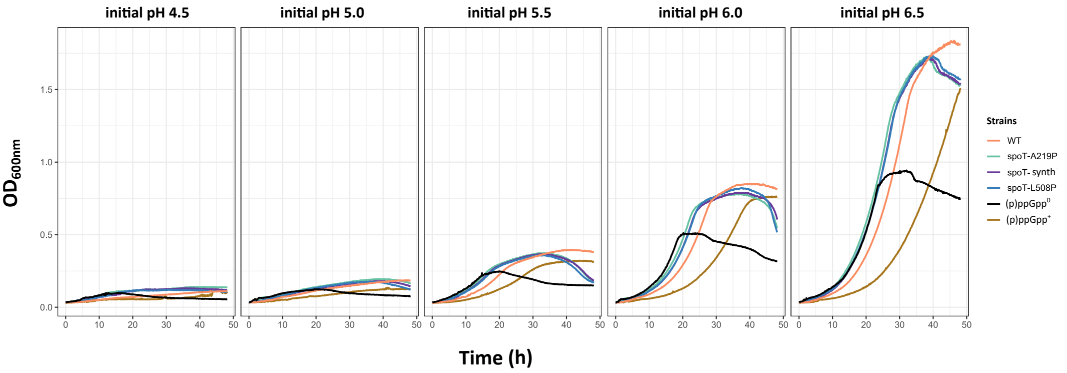


**Figure S5. Growth of *R. pseudosolanacearum* GMI1000 mutants harboring different (p)ppGpp levels at different pH**
Strains were grown in minimal medium supplemented with 56 mM D-glucose. The initial pH of the medium was adjusted using KOH. Data are from one representative experiment, repeated twice independently.

**ALT Text:** Multi-panel line graph showing $OD_{600}$ growth curves over time (0–48 h) for strains with different (p)ppGpp levels across five pH conditions (4.5, 5, 5.5, 6, 6.5). Each panel corresponds to a pH and displays multiple colored lines representing wild-type, *spoT* mutants, and a (p)ppGpp$^{+}$ strain. Growth is similar for all strains. It is minimal at low pH, increases progressively from pH 5.5 to 6.5, and is highest at pH 6.5. The (p)ppGpp$^{\mathbf{0}}$ strain consistently shows reduced maximal optical density at 600 nm compared to other strains.

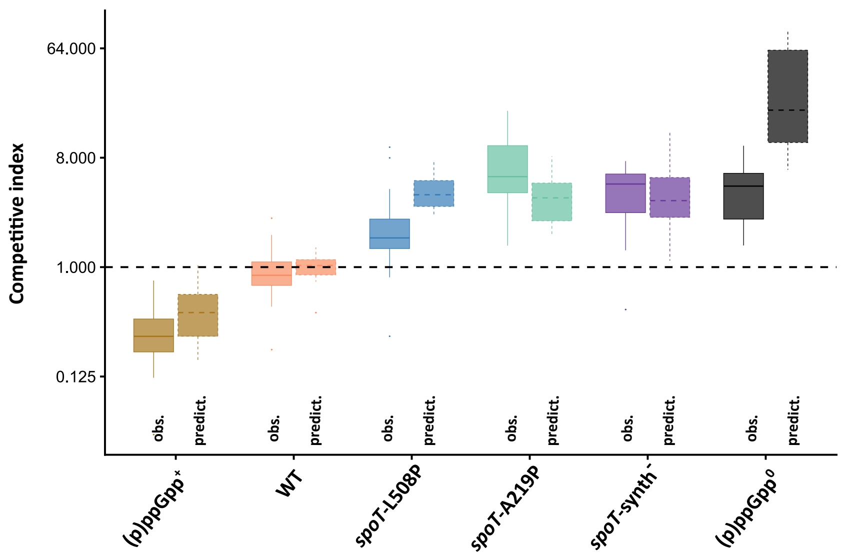


**Figure S6. Comparison of predicted and observed competitive indexes (CIs) of the *R. pseudosolanacearum* GMI1000 *spoT* and (p)ppGpp$^{+}$ mutants in tomato xylem.**
Predictions were calculated using the growth rates measured in minimal medium with L-glutamine as the sole carbon source (raw data are from supplementary table S1E) and an estimated 13 bacterial generations inside the tomato stem (Guidot et al. 2014). obs., observed CIs (data from Fig. 2A). predict., predicted CIs.

**ALT Text:** Two boxplot comparing observed and predicted competitive indexes values in tomato xylem for each strain. Strains include the (p)ppGpp$^{+}$, wild type (GMI1000), three *spoT* mutants (A219P, L508P, E333Q), and the (p)ppGpp$^{0}$ strain. The y-axis in log scale represents CI, with a dashed horizontal line at CI = 1 indicating equal competitiveness. The (p)ppGpp$^{+}$ strain shows CI values below 1, while the wild type is centered around 1. *spoT* mutants display increased CI values above 1, with variability between mutants. The (p)ppGpp$^{0}$ strain shows the highest CI values. Predicted and observed values are generally consistent in their trends across strains.
